# A positively selected microRNA controls a reversible aging program in striated muscle

**DOI:** 10.64898/2026.09.03.749250

**Authors:** Melissa A. Boldridge, Lei Xu, Xiaoyin Wang, Michael Stirm, Gracia Bonilla, Chi Zhu, Justin Y. Lee, Rachelle L. Stark, Sneha Damal Villivalam, Marianne Bengtson Løvendorf, Andreas Petri, Alexandre Wagschal, Caslin Gilroy, Federico Gonzalez, Lei Cai, Peter I. Hecker, Mogens Vyberg, Rahul Almeida, Chaitanya Punnati, Christopher Jin, Theresia M. Schnurr, Dominique O. Riddell, John C. W. Hildyard, Bachuki Shashikadze, Andreas Lange, Nikolai Klymiuk, Thomas Fröhlich, Sakari Kauppinen, Richard J. Piercy, Joshua W. Knowles, Alexander Fay, Sona Kang, Ryan E. Temel, Ruslan I. Sadreyev, Eckhard Wolf, Matthew L. Springer, Anders M. Näär

**Author notes:** These authors contributed equally (Melissa A. Boldridge, Lei Xu, Xiaoyin Wang). Elenae Therapeutics, Inc. (L.X.), Washington University School of Medicine (R.A.), Medical College of Wisconsin (C.J.), Novo Nordisk (T.M.S.). Deceased June 2026. Dr. Springer contributed to this work before his death. Correspondence to Anders M. Näär.

## Abstract

Aging is the primary risk factor for most chronic diseases and is characterized in striated muscle by progressive functional decline, mitochondrial dysfunction, and chronic inflammation. The miR-128-1 locus resides within a positively selected haplotype on chromosome 2q21.3 associated with variation in grip strength, pulmonary function, and cardiometabolic traits in humans. Here, we show that antisense oligonucleotide-mediated inhibition of miR-128-3p restores muscle mass and function in aged mice, improves cardiac function while limiting adverse remodeling following myocardial infarction, and ameliorates skeletal and cardiac muscle pathology in mouse and pig models of Duchenne muscular dystrophy. Across these contexts, miR-128-3p inhibition induces a conserved transcriptional response characterized by activation of mitochondrial programs and suppression of inflammatory and fibrotic signaling, resembling the effects of established longevity interventions. These findings identify miR-128-3p as a regulator of a conserved aging-associated program and establish its inhibition as a strategy to restore tissue function across aging-related muscle pathologies.

**Graphical Abstract:** 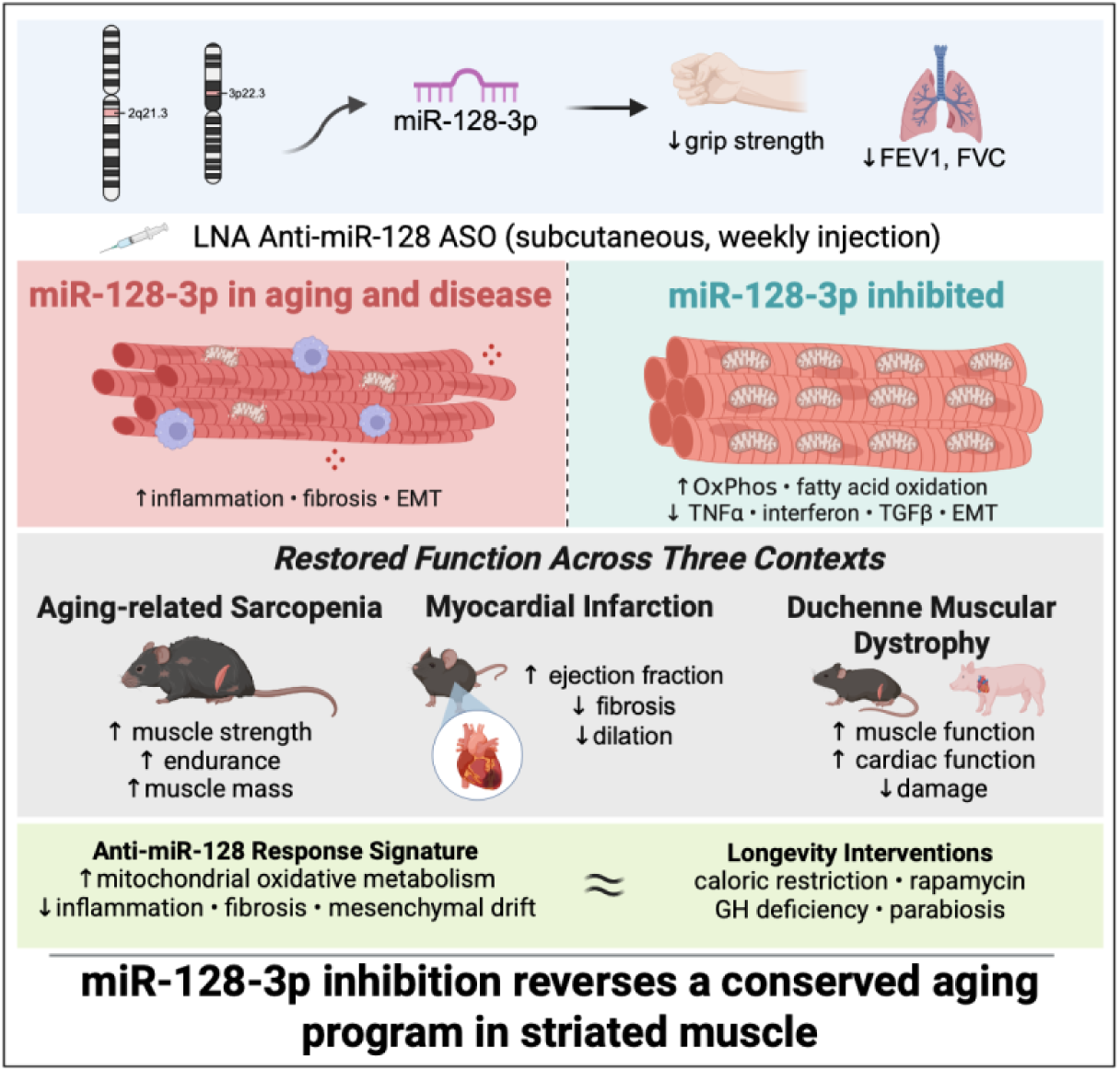

**Highlights:**

- miR-128 loci associate with reduced grip strength; miR-128-1 also with lung function.
- miR-128-3p drives mitochondrial dysfunction and inflammation in striated muscle.
- Anti-miR-128 ASO rescues function in aged, infarcted, and dystrophic muscle.
- Inhibition recapitulates transcriptional effects of longevity interventions.

## Introduction

Two of the strongest predictors of human longevity, grip strength and VO_2_ max, are direct measures of striated muscle performance, reflecting skeletal muscle strength and the integrated function of the cardiovascular and respiratory systems.^1–7^ With advancing age, skeletal muscle mass, quality, and function progressively decline, and accelerated loss of muscle function, termed sarcopenia, is associated with falls, cardiopulmonary disease, impaired recovery from illness, and increased mortality.^8^ At the cellular level, aging muscle is characterized by mitochondrial dysfunction and chronic inflammation, two interconnected processes that reinforce one another.^9^ Consistent with these observations, cross-species transcriptomic analyses have identified a conserved aging signature defined by repression of mitochondrial metabolic pathways and activation of inflammatory and immune responses, a program reversed by longevity interventions including caloric restriction, rapamycin treatment, growth hormone deficiency, and heterochronic parabiosis.^10–12^

This state of mitochondrial dysfunction and chronic inflammation is not unique to physiological aging but is also a prominent feature of striated muscle diseases arising from distinct initiating insults. In Duchenne muscular dystrophy (DMD), loss of dystrophin destabilizes the sarcolemma, rendering muscle fibers susceptible to contraction-induced injury, calcium influx, mitochondrial dysfunction, chronic inflammation, and progressive fibrotic and fatty replacement of contractile tissue.^13–16^ In myocardial infarction (MI), ischemia impairs mitochondrial oxidative phosphorylation, while reperfusion exacerbates injury through reactive oxygen species generation and activation of innate immunity.^17–19^ When this response fails to resolve, persistent inflammation drives adverse remodeling, fibrosis, and progression to heart failure.^18,19^ Thus, natural aging, inherited muscle degeneration, and acute cardiac injury converge on a shared tissue state characterized by impaired mitochondrial function, persistent inflammation, fibrosis, and progressive loss of muscle function.

These observations suggest that physiological aging, acute injury, and genetic disease are driven, at least in part, by a shared molecular program of tissue degeneration. However, the upstream regulators that coordinate this conserved state of mitochondrial dysfunction and chronic inflammation across tissues and disease contexts remain poorly understood. MicroRNAs are well suited to coordinate such programs because they post-transcriptionally regulate networks of target mRNAs, enabling broad control of complex biological states rather than individual signaling pathways.^20,21^

Evolutionary selection can preserve molecular programs that enhance survival under conditions of resource scarcity but become maladaptive with aging and chronic disease. The positively selected haplotype on human chromosome 2q21.3, among the strongest signatures of positive selection in the European genome, has been primarily attributed to selection for lactase persistence. However, this locus is also associated with obesity, type 2 diabetes, and body composition traits, and shows evidence of selection on metabolic phenotypes across multiple mammalian species.^22–25^ We previously identified miR-128-3p, encoded by miR-128-1 located at the center of this locus within the *R3HDM1* gene, as a regulator of circulating lipid levels through genome-wide association analysis of more than 188,000 individuals, and demonstrated that it regulates cholesterol handling and metabolic homeostasis in human cells and mouse metabolic disease models through direct targeting of cholesterol/lipid trafficking genes such as *LDLR* and *ABCA1*.^26^ Our subsequent work showed that miR-128-3p also controls energy expenditure and brown adipocyte function, and that its inhibition ameliorates diet-induced obesity, glucose intolerance, and fatty liver in mice.^25^ These findings established miR-128-3p as a metabolic microRNA whose activity promotes energy conservation, raising the possibility that this pathway may also regulate the transition from adaptive stress resistance to maladaptive tissue aging.

Additional observations support a role for miR-128-3p, the mature product of the miR-128-1 gene, in diseases of striated muscle. miR-128-3p is induced in the heart following MI, and cardiomyocyte-selective genetic deletion or AAV-mediated suppression improves cardiac outcomes after injury.^27,28^ miR-128-3p is also elevated in dystrophic muscle and serum, including in Sapje zebrafish, *mdx* mouse strains, and patients with DMD.^29–31^ Together with its established role in metabolic control, these observations suggested that miR-128-3p might act as a post-transcriptional regulator of a conserved stress program engaged across muscle aging and disease.

Here, we show that genetic variants (single nucleotide polymorphisms; SNPs) at the miR-128-1 locus associate with grip strength and pulmonary function in humans, while SNPs at the miR-128-2 locus associate with weak grip strength and metabolic frailty, together implicating both paralogs in striated muscle function and aging. Using locked nucleic acid (LNA)-modified antisense oligonucleotides (ASO) as molecular tools and potential therapeutic agents, we find that inhibition of miR-128-3p restores muscle function and mass in aged mice, improves cardiac function and limits adverse remodeling following MI in two mouse models, and mitigates skeletal and cardiac muscle pathology in models of DMD across species, including a large mammal. Across these diverse contexts, miR-128-3p inhibition induces a conserved transcriptional response defined by activation of mitochondrial metabolic programs and suppression of inflammatory and fibrotic signaling. Together, these findings identify miR-128-3p as a genetically anchored regulator of a conserved mitochondrial dysfunction-inflammatory axis that underlies striated muscle aging and disease.

## Results

### Genetic Variation at the miR-128 Loci Associates with Age-Related Functional Decline

Mature miR-128-3p is derived from two separate genes, miR-128-1 on chromosome 2, located within *R3HDM1*, and miR-128-2 on chromosome 3, located within *ARPP21*. The two genes show different expression patterns with miR-128-1 driving expression in peripheral tissues whereas miR-128-2 drives expression mainly in the central nervous system. Our previous work links the miR-128-1 locus to metabolic function. GWAS meta-analysis of more than 188,000 individuals identified the miR-128-1 locus among 69 microRNA loci associated with circulating lipid levels, with expression QTL analyses linking genetic variation directly to miR-128-3p expression in metabolically active tissues.^26^ An independent population genetics analysis shows that this locus exhibits one of the strongest signals of recent positive selection in the European genome, consistent with selection acting on pathways governing energy balance and metabolic efficiency.^25^ Given that metabolic dysfunction underlies many aging phenotypes, and that this locus had been linked to metabolic traits, we hypothesized that genetic variation at the miR-128-1 and -2 loci might influence human aging trajectories.

To characterize the phenotypic landscape associated with the miR-128 loci, we used OpenTargets to investigate phenotypes with fine-mapped credible sets overlapping with miR-128 loci. We highlight rs1446585, rs1446584, and rs1438307 in the miR-128-1 locus and rs1513475 in the miR-128-2 locus, which are members of fine-mapped credible sets. Consistent with our previous work,^26^ the miR-128-1 SNPs lie within credible sets associated with measures of cholesterol including total cholesterol, non-HDL cholesterol (rs1446585), ApoB (rs1446585), and LDL (rs1446585) (Fig. 1A, S1A-B). All three miR-128-1 SNPs also associate with measures of body composition and adiposity including BMI, waist-hip ratio, body fat %, morbid obesity, and obesity (rs1446585) (Fig. 1A, S1A-B). This aligns with our previous studies showing miR-128-3p inhibition prevents diet-induced obesity in mice.^25^ Our current analyses of SNP rs1513475 also show this association with body composition and adiposity related traits in the miR-128-2 locus that have not previously been described (Fig. 1B).

**Figure 1.**
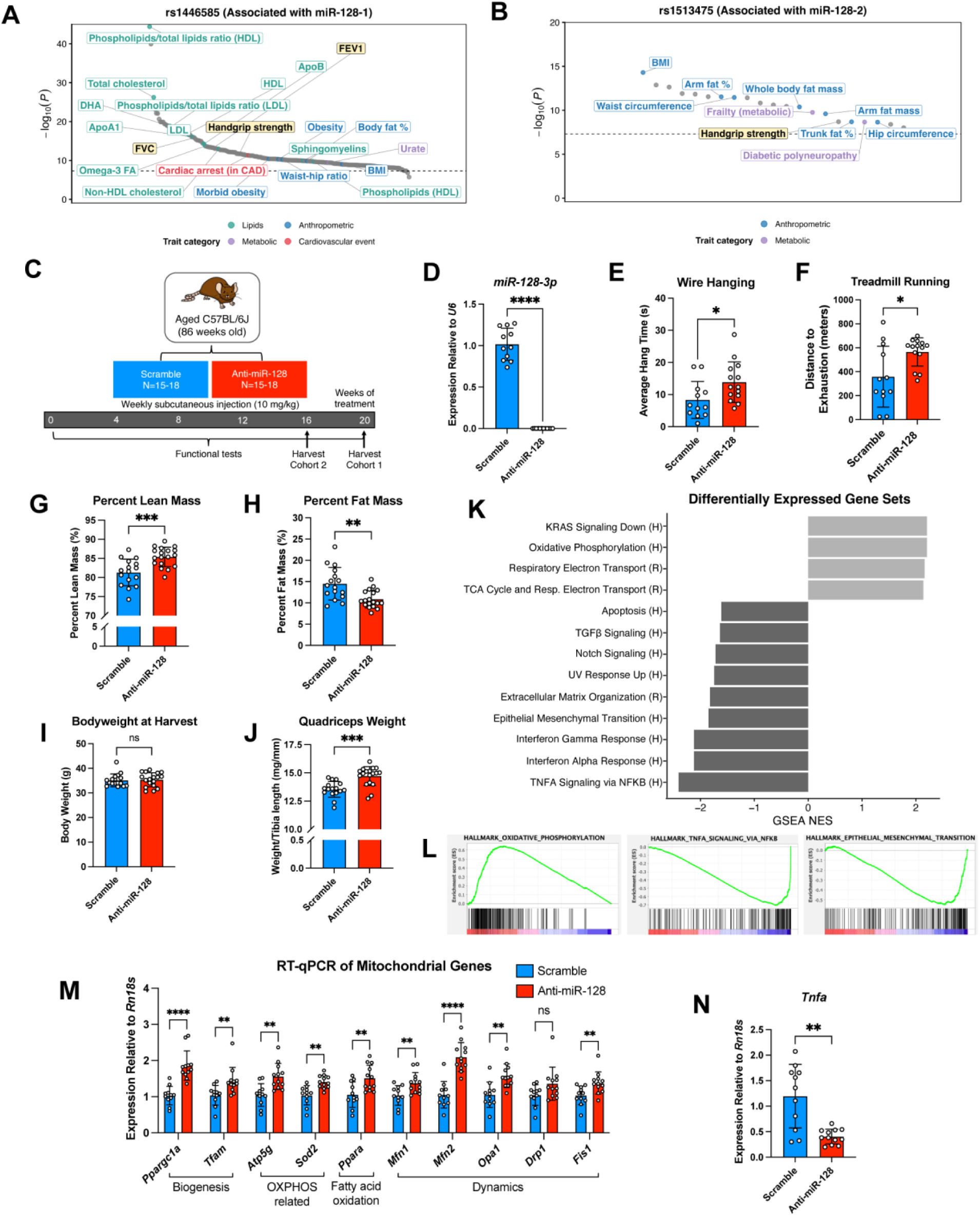
SNPs in the miR-128 loci are associated with muscle and pulmonary function in humans and anti-miR-128 improves muscle function and transcriptional programs in aged mice. (A) PheWAS plot of the single nucleotide polymorphism (SNP) rs1446585 at the miR-128-1 locus. Dashed line denotes genome-wide significance (P<5 x 10^-8^). (B) PheWAS plot of the single nucleotide polymorphism (SNP) rs1513475 at the miR-128-2 locus. Dashed line denotes genome-wide significance (P<5 x 10^-8^). (C) Experimental design diagram. 86-week-old mice were treated with anti-miR-128 or scramble once weekly and analyzed after 20 weeks (cohort 1) or 16 weeks (cohort 2) of treatment. Muscle function was assessed longitudinally. (D) *miR-128-3p* expression relative to *U6* in gastrocnemius muscle. (E) Forepaw wire hanging performance at 15 weeks of treatment from Cohort 1. (F) Endurance treadmill running until exhaustion at 13 weeks of treatment from Cohort 1. (G-H) Body composition measured by EchoMRI. Percent lean mass (G) and fat mass (H) at 15 weeks of treatment from cohort 2. (I) Body weight at harvest from cohort 2. (J) Quadriceps muscle wet tissue weight normalized to tibia length from cohort 2. (K) Functional category enrichment in upregulated genes (light gray) and down-regulated genes (dark gray) by anti-miR-128 compared with scramble treated aged mouse gastrocnemius muscle from cohort 1. Bars represent normalized enrichment score (NES) according to Gene Set Enrichment Analysis (GSEA). (H, Hallmark; R, Reactome; FDR<0.05) (L) GSEA enrichment plots of the Oxidative Phosphorylation Hallmark gene set (left), the TNFα signaling via NFκB Hallmark gene set (middle), and the Epithelial Mesenchymal Transition Hallmark gene set (right). (M) RT-qPCR of GA muscle of mitochondrial biogenesis, oxidative phosphorylation, fatty acid oxidation, and dynamics related genes from cohort 1. (N) RT-qPCR of GA muscle of *Tnfa* expression from cohort 1. Data are presented as mean +/- SD. N=11-12 (D, K-N); N=12-14 (E-F); N=16-18 (G-J). ns, not significant, *p<0.05, **p<0.01,***p<0.001,****p<0.0001 (unpaired, two-tailed t-test).

In addition to the metabolic traits anticipated from our previous investigations, all examined SNPs in the miR-128-1 and miR-128-2 loci lie within credible sets for hand grip strength, one of the most robust biomarkers of biological aging and all-cause mortality (Fig. 1A, S1A-B).^1–3^ The miR-128-1 SNPs associate with markers of pulmonary function as well, specifically forced vital capacity (FVC; rs1446585) and forced expiratory volume in one second (FEV1) (Fig. 1A, S1A-B). Together, these traits converge on striated muscle function, providing support for a potential role of miR-128-3p in regulating programs driving muscle aging.

### Antisense Targeting of miR-128-3p Provides Robust and Precise Inhibition with Favorable Tolerability Profile

To examine the role of miR-128-3p in muscle aging, we have employed ASOs as both a useful molecular tool and a potential therapeutic agent. We have designed an LNA-modified ASO targeting miR-128-3p, hereafter referred to as anti-miR-128, which can be delivered by subcutaneous injection in saline. Subcutaneous administration of anti-miR-128 in C57BL/6J mice provides robust knockdown (71-93%) in the liver at doses ranging from 1.25-10 mg/kg body weight compared with a scramble control ASO (scramble) after 28 days (Fig. S2A). Cardiac knockdown can also be achieved in this dosing range with nearly complete knockdown at 5-10 mg/kg (Fig. S2B). In skeletal muscle (a generally difficult to target tissue), near complete knockdown is also achieved with 10 mg/kg dosing (Fig. S2C). Using this dose of 10 mg/kg, we have also shown that miR-27a-3p, which shares an 8 nucleotide overlap with miR-128-3p and has functions in metabolism,^32,33^ maintains the same expression level in skeletal muscle compared to muscle of scramble ASO treated mice (Fig. S2D), demonstrating high specificity of our anti-miR-128 ASO. Thus, 10 mg/kg dosing was adopted for future rodent studies.

Kinetic analysis of anti-miR-128 following a single dose revealed tissue-specific patterns of uptake and clearance. Anti-miR-128 reached liver, kidney, heart, and skeletal muscle within 24 hours and was cleared from the serum within 10 days (Fig. S2E-J). Notably, cardiac and skeletal muscle retained high anti-miR-128 levels throughout the 10-day period. These pharmacokinetic properties informed a weekly dosing schedule to maintain effective skeletal and cardiac muscle knockdown in subsequent studies.

Anti-miR-128 was well tolerated with prolonged administration in both mice and cynomolgus monkeys. When delivered weekly at 10 mg/kg to aged rodents over 20 weeks, no adverse changes to liver or kidney histology were observed (Fig. S3 A-B) and serum ALT and AST activity, blood urea nitrogen, and creatinine levels remained unchanged compared with scramble treated animals (Fig. S3 C-F). Male cynomolgus monkeys fed with a fast-food diet with high fructose drink to induce metabolic syndrome were treated with anti-miR-128 at 5 mg/kg (n=4) or saline (n=3) by weekly subcutaneous injection for 24 weeks, with serum chemistry and hematology assessed at 14 timepoints. Serum markers of hepatic and renal function, including ALT and AST (Fig. S3G-H; Table 1) and BUN and creatinine (Fig. S3I-J; Table 1) respectively, remained stable throughout the study and did not show significant differences between treatment groups at any of the measured timepoints. Histological examination of liver and kidney tissue at the study endpoint revealed no treatment-related pathological changes (Fig. S3K-L). Platelet counts, the principal class liability for phosphorothioate ASO backbone chemistry,^34^ rose over the dosing period in treated animals (291 to 396 K/µL) as they did in controls (368 to 433 K/µL), and no treated animal fell below 191 K/µL at any of fourteen draws. These data support a favorable tolerability profile for anti-miR-128 in mice and non-human primates.

### Anti-miR-128-3p Improves Skeletal Muscle Mass, Performance, and Transcriptional Programs in Aged Mice

Having established genetic links between miR-128-3p and human aging phenotypes as well as the dosing and safety of chronic anti-miR-128 administration, we asked whether miR-128-3p activity drives age-related muscle dysfunction and whether this could be therapeutically reversed. To test this, we treated 86-week-old C57BL/6J mice with anti-miR-128 or scramble at 10 mg/kg weekly for 16 or 20 weeks (Fig. 1C). During these studies, 4 scramble and 3 anti-miR-128 treated mice in the 20-week cohort (cohort 1), and 2 scramble treated mice in the 16-week cohort (cohort 2), died prior to the study endpoint. Anti-miR-128 treatment robustly suppressed miR-128-3p expression in gastrocnemius muscle (Fig. 1D). We performed muscle function tests throughout the study to assess the impact of anti-miR-128 treatment on aging-related functional decline. Passive wire hanging revealed a significant increase in average hang time by 66% with anti-miR-128 compared to scramble at 15 weeks of treatment (Fig. 1E). Tests at 5 and 9 weeks of treatment similarly showed a trending increase in average hang time (Fig. S4A-B). Treadmill run time to exhaustion was also significantly increased by 58% with anti-miR-128 at 13 weeks of treatment using a low-intensity treadmill running protocol (Fig. 1F). A high intensity running protocol showed a significant, though somewhat less robust, increase (41% compared to scramble) at 8 weeks of treatment (Fig. S4C). Increases in treadmill run time with anti-miR-128 were also observed in a second independent cohort at 5 and 12 weeks of treatment (Fig. S4D-E). These data suggest that anti-miR-128 can mitigate the loss of muscle function that is characteristic of aging in mice.

Anti-miR-128 treatment not only improved functional performance but also increased muscle mass. EchoMRI at 15 weeks of treatment showed a significant increase in lean body mass with anti-miR-128, 81.4% to 85.4% (Fig. 1G), accompanied by a corresponding significant reduction in fat mass from 14.5% to 10.9% (Fig. 1H), and no change in body mass (Fig. 1I). This is consistent with our previous observations that miR-128-3p inhibition reduces fat mass in a diet-induced obesity mouse model.^25^ At harvest, quadriceps weight (normalized to tibia length) was significantly higher with anti-miR-128 treatment (Fig. 1J). Additionally, the distribution of muscle fiber cross sectional area (CSA) in TA muscle showed a significant shift toward larger fibers with anti-miR-128 treatment (Fig. S4F-H). Thus, anti-miR-128 treatment is associated with both increased skeletal muscle mass and functional improvement.

To define the molecular basis for improved muscle performance and mass, we performed bulk RNA sequencing on gastrocnemius muscle from aged mice treated with anti-miR-128 or scramble for 20 weeks. Gene set enrichment analysis (GSEA) revealed upregulation of mitochondrial metabolism pathways (Fig. 1K) including oxidative phosphorylation (Fig. 1L, left) and the TCA cycle. At the individual gene level, RT-qPCR analysis revealed significant induction of regulators of mitochondrial biogenesis (*Ppargc1a, Tfam*), oxidative phosphorylation components (*Atp5g, Sod2*), mitochondrial fatty acid oxidation (*Ppara*), and genes involved in mitochondrial dynamics (*Mfn1, Mfn2, Opa1, Fis1*) (Fig. 1M). This shows de-repression of validated miR-128-3p targets (*Ppargc1a*, *Ppara*, *Tfam*, and *Opa1*) as well as increased expression of other mediators of mitochondrial function that may be indirect targets. In contrast, GSEA revealed downregulation of several gene sets related to inflammation and fibrosis, including Interferon alpha and gamma responses, TNFα signaling via NFκB (Fig. 1L, middle), TGF*β* signaling, and genes associated with epithelial mesenchymal transition (EMT; Fig. 1L, right) (Fig. 1K). Downregulation of *Tnfa* expression was directly confirmed by RT-qPCR (Fig. 1N). These reciprocal changes in mitochondrial and inflammatory programs are consistent with a model in which miR-128-3p-mediated mitochondrial dysfunction contributes to inflammatory activation in aged muscle, and that miR-128-3p inhibition restores mitochondrial programs while attenuating chronic inflammation associated with aging.

### Inhibition of miR-128-3p Mitigates Adverse Cardiac Remodeling Following Myocardial Infarction

Encouraged by the beneficial effects of anti-miR-128 in natural aging, we next asked whether miR-128-3p inhibition could also confer benefit in aging-associated diseases of striated muscle. We focused on the heart because of its exceptional dependence on mitochondrial oxidative metabolism to meet its high energetic demands. Previous studies have implicated miR-128-3p in mediating adverse remodeling following myocardial infarction.^27,28^ Thus, we examined the therapeutic efficacy of anti-miR-128 in a mouse ischemia reperfusion (I/R) MI model generated by 30 minutes of left anterior descending (LAD) coronary artery occlusion (Fig. 2A). In this model, weekly administration of anti-miR-128 (10 mg/kg) significantly improved left-ventricular ejection fraction (EF) measured by serial echocardiography over 28 days (Fig. 2B). While scramble-treated mice exhibited a reduction in EF from 55.2% before MI to 33.6% 28 days later, anti-miR-128 treated mice with a similar baseline EF rebounded to 51.8% by 28 days post-I/R. Treatment also attenuated adverse ventricular remodeling post-MI, including reduced left ventricular dilation, reflected in reduced end-systolic volume (ESV) and end-diastolic volume (EDV) respectively (Fig. 2C-D). Notably, a single dose of anti-miR-128 given either during ischemia (Fig. 2B-D) or 2 hours after reperfusion, a more therapeutically relevant timepoint, (Fig. 2E-G) produced comparable improvements in EF, ESV, and EDV. These data demonstrate the therapeutic efficacy of anti-miR-128 in mitigating adverse remodeling and restoring cardiac function in mouse models of MI with clinically relevant delivery timepoints.

**Figure 2.**
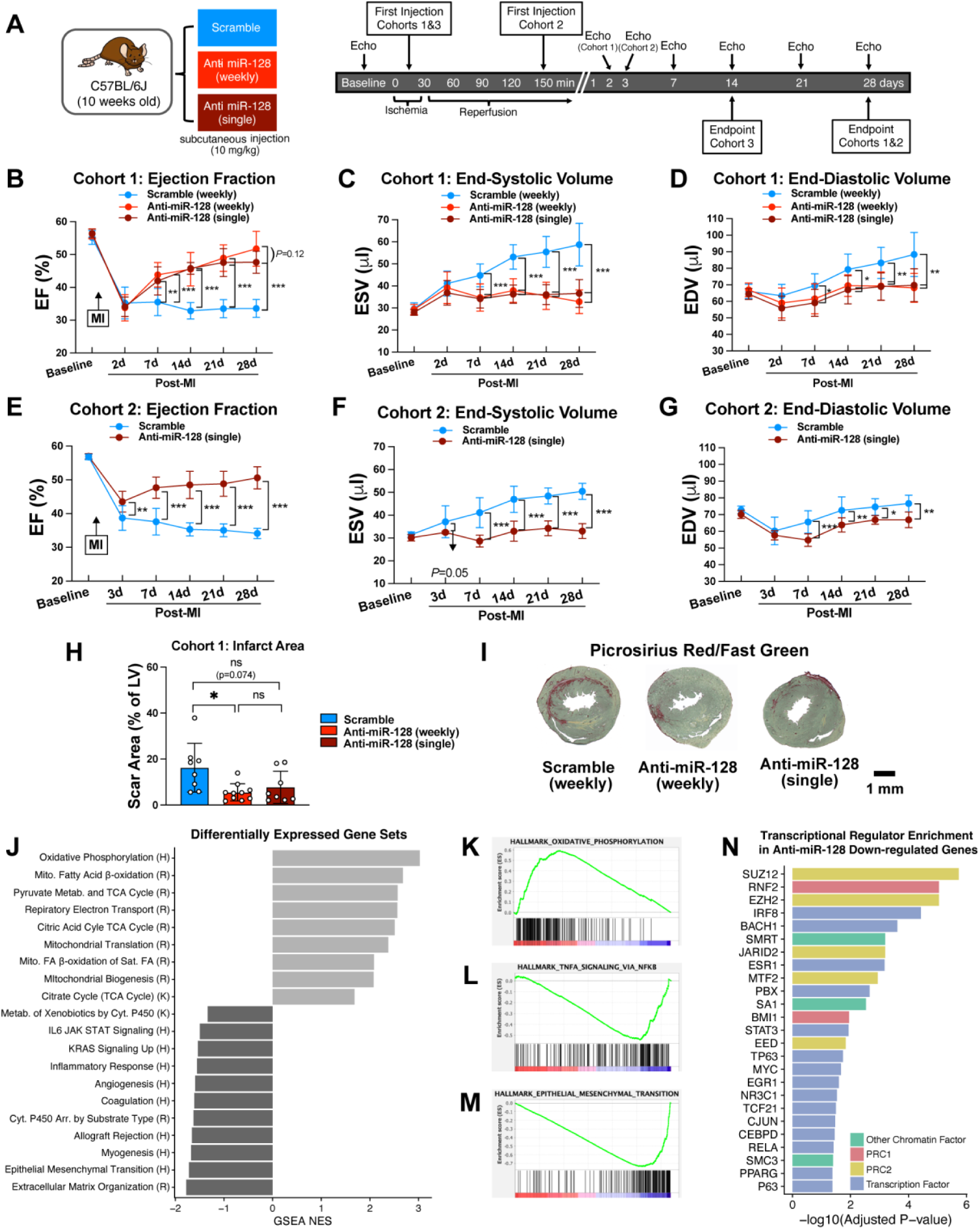
Anti-miR-128 improves cardiac function, reduces fibrosis, and suppresses inflammatory and profibrotic gene expression programs following MI. (A) Experimental timeline. Mice underwent ischemia-reperfusion (I/R) MI and received anti-miR-128 or scramble, with echocardiography and tissue collection at the indicated timepoints. Anti-miR-128 or scramble was delivered at 10 mg/kg as either a single injection after 15 min of ischemia (cohorts 1, single and cohort 3), a similarly timed initial injection followed by weekly injections for 3 weeks (cohort 1, weekly), or a single injection 2 hours after the onset of reperfusion (cohort 2). (B-D) Serial echocardiographic measurements of left ventricular (LV) ejection fraction (EF) (B), end-systolic volume (ESV) (C), and end-diastolic volume (EDV) (D) over 28 days post-I/R in cohort 1. Mice received weekly scramble, weekly anti-miR-128, or a single injection of anti-miR-128 15 minutes after the onset of ischemia. (E-G) Serial echocardiographic measurements of LV EF (E), ESV (F), and EDV (G) over 28 days post-IR in cohort 2. Mice received a single injection of anti-miR-128 or scramble two hours after reperfusion. (H) Infarct scar area quantified 28 days post-I/R by Picrosirius Red/Fast Green staining. (I) Representative myocardial sections from each of the three groups in (H) showing the section with the largest scar area from each heart, together with the mean infarct scar area for each group. (J) Functional category enrichment in upregulated genes (light gray) and down-regulated genes (dark gray) by anti-miR-128 compared with scramble treated mouse hearts 14 days post-I/R MI (cohort 3). Bars represent normalized enrichment score (NES) according to Gene Set Enrichment Analysis (GSEA). (H, Hallmark; R, Reactome; FDR<0.05) (K-M) GSEA enrichment plots for the Oxidative Phosphorylation Hallmark gene set (K), TNFα signaling via NFκB Hallmark gene set (L), and Epithelial-Mesenchymal Transition Hallmark gene set (M). (N) Enrichment of transcription factor and chromatin regulator binding signatures among 445 genes down-regulated by anti-miR-128 treatment in post-MI cardiac tissue. Bars are colored by category and represent -log10 of the adjusted P-value (FDR) of enrichment according to Enrichr (ChEA database). Only results with FDR < 0.05 are shown. Data are presented as mean +/-SD. N=8-10 per group (B-D); N=5 per group (E-G); N=8-10 (H); N=4 per group (J-M). *p<0.05, **p<0.01,***p<0.001,****p<0.0001 (two-way ANOVA followed by Šidák’s multiple comparisons test (B-G); one-way ANOVA followed by Tukey’s multiple comparisons test (H)).

To better understand the impact of miR-128-3p inhibition on MI-induced injury, we performed fibrosis staining. Histological analysis demonstrated significantly reduced infarct scar area with weekly administration and a trending decrease in infarct scar area with a single dose 28 days post-I/R (Fig. 2H-I; Fig. S5A). To gain a better understanding of myocardial remodeling in response to injury, we examined anterior and posterior wall thickness at the apical infarct zone and the basal remote zone by echocardiography. In the scramble-treated group, wall thickness at the apical infarct zone, both anteriorly and posteriorly, was decreased 28 days post-I/R compared to baseline (Fig. S5B). Anti-miR-128 treatment mitigated this decrease, suggesting prevention of injury-associated damage to the myocardium. At the basal remote zone, wall thickness was increased in scramble-treated mice, suggesting adverse hypertrophic remodeling post-MI (Fig. S5C). In contrast, anti-miR-128 treatment attenuated wall thickening at the basal remote zone, particularly in the posterior wall (Fig. S5C, right). Together, these data show that anti-miR-128 prevented wall thinning at the damaged apical infarct zone as well as compensatory wall hypertrophy at the basal remote zone, ameliorating adverse left-ventricular remodeling post-I/R.

The heart relies predominantly on mitochondrial oxidative phosphorylation to generate ATP and meet its high energetic demands. Myocardial ischemia disrupts mitochondrial metabolism and necessitates a shift toward glycolysis, and subsequent reperfusion further exacerbates mitochondrial injury and inflammation.^17,35^ Although inflammation is an essential component of the myocardial injury response, failure to resolve this response drives adverse ventricular remodeling and progression to heart failure.^18,19^ Because anti-miR-128 improved skeletal muscle function during aging, we asked whether it similarly restored conserved transcriptional programs in the post-MI heart. RNA sequencing of left ventricular tissue at 14 days post-I/R revealed that anti-miR-128 treatment (versus scramble) resulted in upregulation of mitochondrial gene expression programs (Fig. 2J), with significant enrichment of the oxidative phosphorylation pathway (Fig. 2K), fatty acid beta-oxidation, TCA cycle, mitochondrial translation, and mitochondrial biogenesis. These changes were accompanied by suppression of inflammatory pathways (Fig. 2J), including inflammatory response, IL6/JAK/STAT signaling, and TNFα signaling via NFκB (Fig. 2L), as well as EMT-associated gene programs (Fig. 2M). We next asked which regulators might account for this downregulated gene signature. Genes downregulated by anti-miR-128 were significantly enriched in gene sets for targets of Polycomb repressive complex components including PRC2 subunits SUZ12, EZH2, JARID2, MTF2, and EED as well as PRC1 subunits RNF2 and BMI1 (Fig. 2N). Notably, SUZ12 is a direct target of miR-128-3p suggesting that anti-miR-128 enhances PRC2 mediated transcriptional repression.^27^ These findings support a conserved role for miR-128-3p in mediating damage across striated muscle diseases associated with aging and suggest shared nodes of action via mitochondrial dysfunction and chronic inflammation.

The I/R MI model recapitulates the reperfusion injury that occurs following restoration of LAD coronary blood flow. However, because mice exhibit substantial tolerance to myocardial injury, large infarcts are often required to model progressive heart failure.^36^ We therefore evaluated anti-miR-128 in a complementary model of permanent ligation of the LAD, which produces sustained ischemic injury. Anti-miR-128 treatment similarly improved cardiac function and attenuated adverse remodeling in this model (Fig. 3A-C). Specifically, anti-miR-128 treatment improved left ventricular EF and reduced ESV, although EDV was not significantly affected (Fig. 3A). Although infarct size was unchanged using this model, fibrosis at the scar zone was reduced with treatment (Fig. 3B-C). Thus, the beneficial effects of miR-128-3p inhibition are not limited to reperfusion injury but extend to sustained ischemic injury and adverse remodeling.

**Figure 3.**
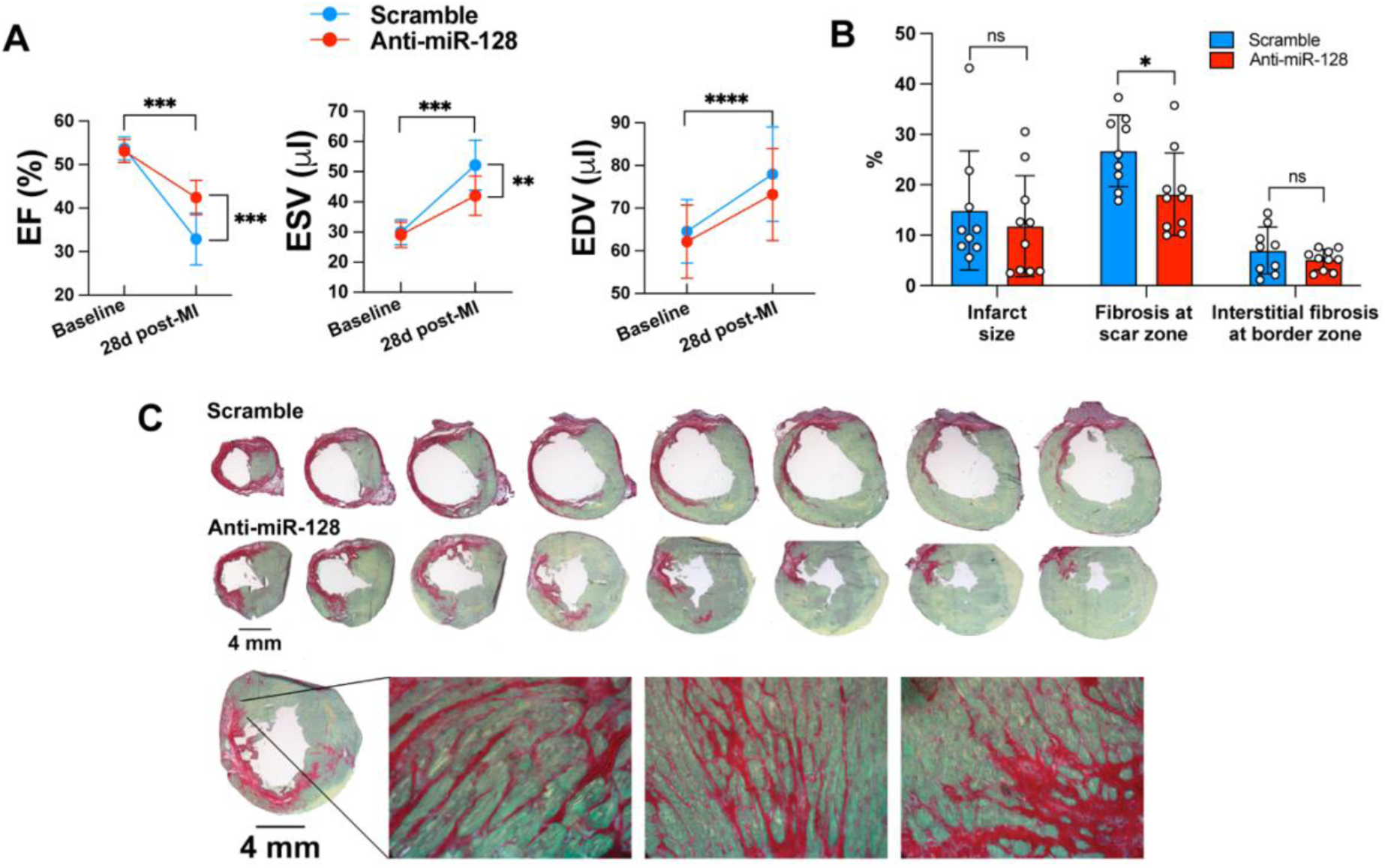
Anti-miR-128 improves cardiac function and reduces fibrosis after permanent ligation MI. (A) Echocardiography from mice treated with weekly injections (10 mg/kg) of scramble or anti-miR-128 post permanent ligation MI. Left ventricular ejection fraction (EF) (left), end-systolic volume (ESV) (middle), and end-diastolic volume (EDV) (right) at baseline and 28 days post-MI. (B) Infarct size, fibrosis at scar zone, and interstitial fibrosis at border zone at 28 days post-permanent ligation MI. (C) Representative serial myocardial sections stained with Picrosirius Red and Fast Green from scramble-treated and anti-miR-128-treated mice after permanent ligation MI. The upper two rows show serial sections to quantify infarct size, and lower row shows representative higher-magnification images used to quantify fibrosis at the scar zone. Heart samples were sectioned transversely at 10 µm thickness with an interval of 300 µm between each section. Eight sections from apex to base of the ventricle were stained and used for infarct scar size measurement. Data are presented as mean +/- SD. N=9-11 per group (A); N=9-10 per group (B). *p<0.05, **p<0.01,***p<0.001,****p<0.0001 (two-way ANOVA followed by the Šidák method (A) and unpaired two-tailed t-test (B)).

### Anti-miR-128 Improves Muscle Function and Attenuates Pathology in Dystrophic Muscle

After observing striking benefit from anti-miR-128 treatment in two aging-associated striated muscle diseases, we wanted to examine its effect in a genetic disease of striated muscle function. Duchenne muscular dystrophy (DMD) patients and animal models exhibit an increase in miR-128-3p expression in serum and skeletal muscle (Fig. 4A-C, E).^29–31^ To determine whether miR-128-3p contributes to a shared stress program in dystrophic muscle, we administered anti-miR-128 at 10 mg/kg to *mdx*^5c^*^v^* mice for 3-6 weeks beginning at 3 weeks of age (Fig. 4D). Anti-miR-128 robustly suppressed miR-128-3p expression in gastrocnemius muscle (Fig. 4E) and reduced circulating creatine kinase levels (Fig. 4F), indicating a reduction of ongoing muscle damage. Muscle function as measured by wire hanging and treadmill endurance was significantly improved compared with scramble-treated littermates (Fig. 4G-H). Histology showed significantly reduced fiber necrosis (Fig. 4I-J) and significantly reduced muscle fibrosis (Fig. 4K-L) with anti-miR-128 treatment. Furthermore, we also observed an increase in oxidative type IIa fibers and a reduction in glycolytic type IIb fibers (Fig. 4M-N). When examining mitochondrial ultrastructure, *mdx*^5c^*^v^*mice showed a clear misalignment of mitochondria along the myofibrils (Fig. 4O, yellow arrows) and perinuclear accumulation (Fig. 4O, white arrows) compared with wildtype controls, which is characteristic of regenerating muscle fibers.^37,38^ These abnormalities were attenuated by anti-miR-128 treatment. The increased oxidative fibers and improvement in mitochondrial ultrastructure was also accompanied by an increase in mitochondrial DNA copy number suggesting greater mitochondrial mass (Fig. 4P). Thus, the observed improvement in muscle function is driven by improved muscle tissue structure including fiber hypertrophy, improved fiber survival, and reduced non-contractile fibrotic tissue. This is accompanied by increased mitochondrial content, oxidative fibers, and improved mitochondrial ultrastructure.

**Figure 4.**
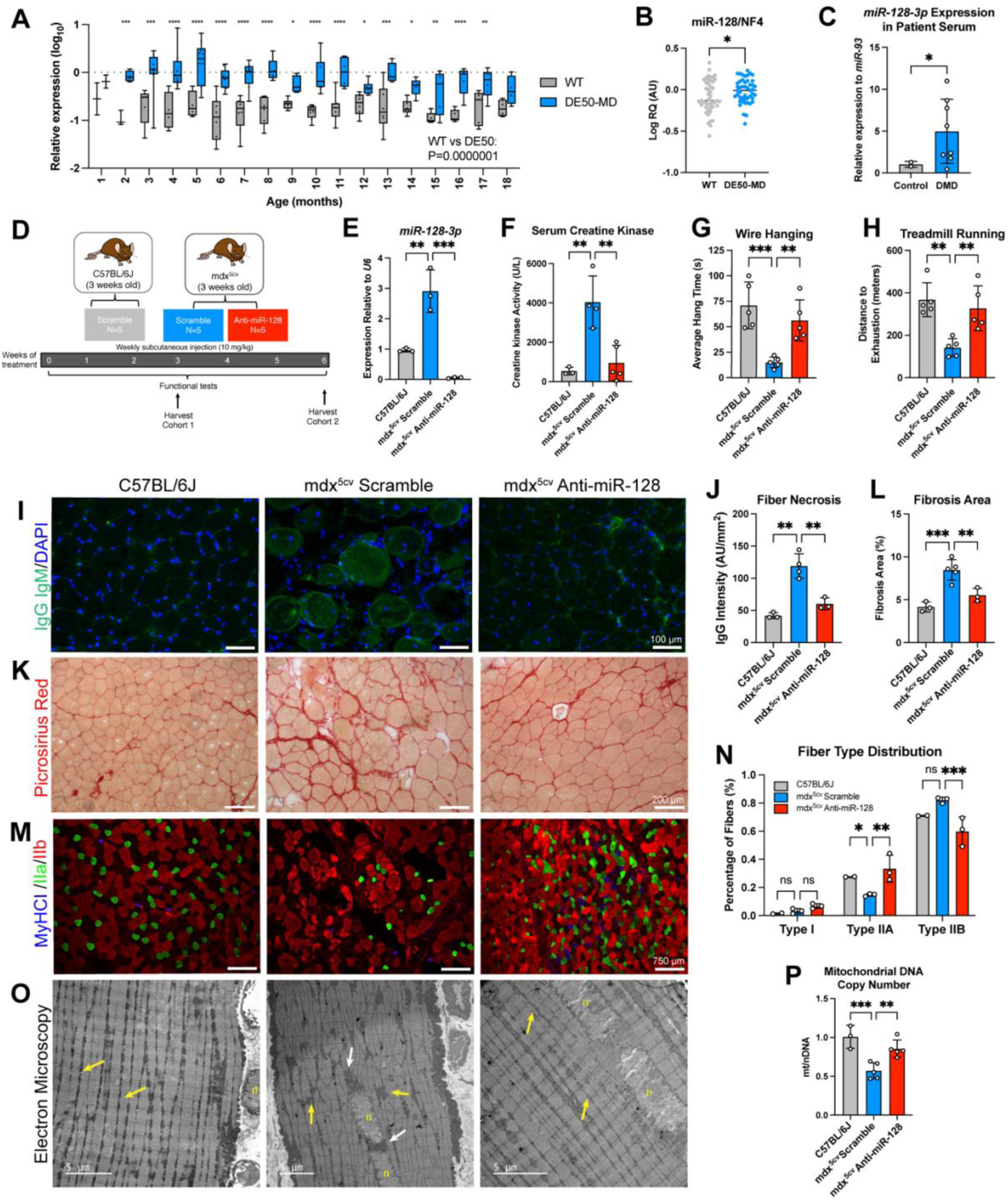
miR-128-3p expression is elevated in DMD dogs and patients and anti-miR-128 improves skeletal muscle function, enhances mitochondrial homeostasis, improves muscle remodeling in the *mdx*^5c^*^v^*mouse. (A) miR-128-3p expression in DE50-MD and WT canine serum samples collected monthly between 1 and 18 months of age, normalized to miR-223 expression. Samples were obtained from N = 8 WT and N = 12 DE50-MD dogs. (B) miR-128-3p expression in DE50-MD and WT canine muscle samples, normalized to a four-microRNA normalization factor (NF4; geometric mean of the relative quantities of miR-191, let-7b, miR-125a and miR-15a). The graph summarizes a genotype comparison by linear mixed model of WT (n = 47) and DE50-MD (n = 47) muscle samples, comprising vastus lateralis sampled longitudinally from N = 6 WT and N = 6 DE50-MD dogs at 3, 6, 9, 12, 15 and 18 months, together with cranial tibial, semimembranosus, triceps and diaphragm samples from N = 3 WT and N = 3 DE50-MD dogs at 18 months. One DE50-MD dog contributed to both sets, giving a total of N = 9 WT and N = 8 DE50-MD unique dogs. (C) Serum miR-128-3p expression relative to miR-93 in patients with DMD and in comparison participants from the same clinic with unrelated neuromuscular diagnoses. (D) Schematic of experimental design. *mdx*^5c^*^v^* mice were treated with anti-miR-128 or scramble at 10 mg/kg weekly for 3 weeks (cohort 1) or 6 weeks (cohort 2) starting from 3 weeks of age and muscle function was assessed throughout the study. (E) *miR-128-3p* expression level relative to *U6* in the gastrocnemius muscle wildtype and *mdx*^5c^*^v^* mice treated with scramble and anti-miR-128. (F) Creatine kinase activity in serum at study endpoint (cohort 1). (G) Forelimb wire hanging at 3 weeks of treatment (cohort 1). (H) Distance run on treadmill until exhaustion at 3 weeks of treatment (cohort 1). (I-J) Representative images of IgG/IgM/DAPI immunostaining of TA muscle from cohort 2 (I) and quantification of necrotic muscle fibers in (J). (K-L) Representative images of Picrosirius Red staining of TA muscle from cohort 2 (K) and quantification of fibrosis in (L). (M-N) Representative images of myosin heavy chain (MyHC) fiber type staining (Type I blue, Type IIa green, Type IIb red) of TA muscle from cohort 2 (M) with quantification of muscle fiber types in (N). (O) Representative electron microscopy images of subsarcolemmal and intermyofibrillar mitochondria in soleus muscle from cohort 1 (n: nucleus). (P) Mitochondrial DNA copy number relative to nuclear DNA quantified by RT-qPCR from soleus muscle from cohort 1. Data are presented as mean +/-SD. N=96 (WT) and 112 (DE50-MD) serum samples from 8 WT and 12 DE50-MD dogs, with N ≥ 3 dogs per genotype per timepoint at all timepoints (mean 5-6) except 1 month (A); from N=9 WT and N=8 DE50-MD dogs (B); N=2-7 per group (C), N=3 per group (E); N=3-5 per group (F, P); N=5 per group (G-H); N=3-4 (J); N=4-5 (L); N=2-3 (N). *p<0.05, **p<0.01, ***p<0.001 (linear mixed model analysis (A-B), unpaired, two-tailed t-test (C), one-way ANOVA followed by Dunnett’s multiple-comparisons test (E-H, J, L, N, P))

To better understand the transcriptional program driving these observed functional and tissue improvements, we conducted bulk RNA sequencing on GA muscle from *mdx*^5c^*^v^*mice treated with anti-miR-128 or scramble for 3 weeks. Transcriptomic analysis revealed enrichment of mitochondrial metabolic pathways, including oxidative phosphorylation, TCA cycle, and pyruvate metabolism (Fig. 5A, B), together with suppression of inflammatory gene programs such as TNFα/NF-κB signaling and interferon responses (Fig. 5A, C), as well as EMT-associated genes (Fig. 5A, D). RT-qPCR supported upregulation of regulators of mitochondrial biogenesis (*Ppargc1a, Tfam*), dynamics (*Mfn1, Mfn2, Opa1, Drp1, Fis1*), metabolic control (*Sod2, Ppara*), and cellular energy sensing and regulation (*Prkaa2, Sirt1*) and suppression of inflammation (*Tnfa*) (Fig. 5E-F). At the protein level, we also observed upregulation of the mitochondrial biogenesis regulator PGC1α, as well as PPARα and CPT1b, which are important for mitochondrial fatty acid oxidation (Fig. 5G-H). Several of these genes, including *Ppargc1a*, *Tfam*, *Opa1*, *Ppara*, and *Sirt1*, are validated direct targets of miR-128-3p, so their induction is consistent with direct de-repression rather than a secondary consequence of improved tissue state alone. ^25,28,39,40^ These gene expression changes resemble those observed in aged muscle and post-MI hearts after anti-miR-128 treatment, suggesting a shared function of miR-128-3p across striated muscle diseases and aging.

**Figure 5.**
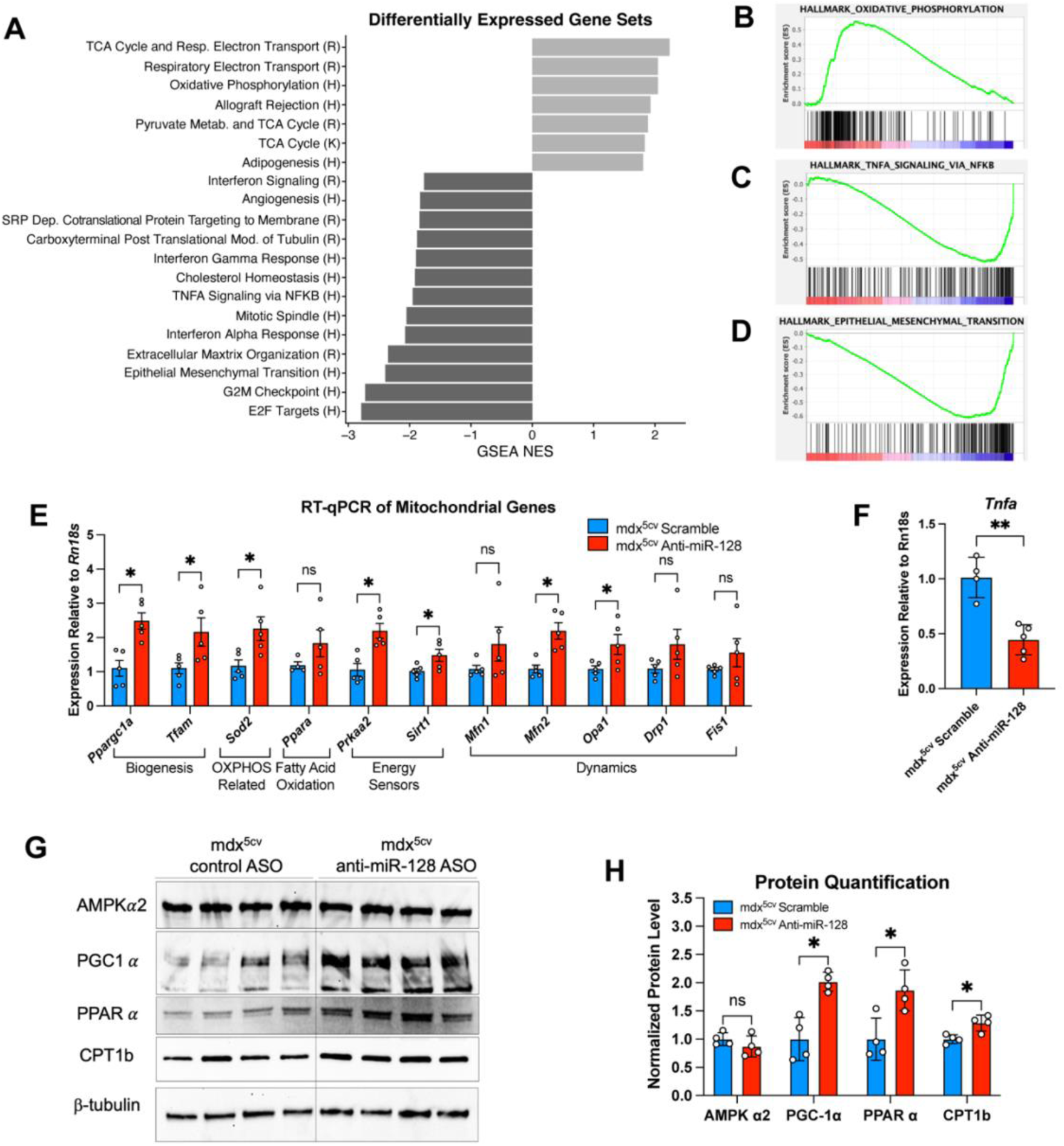
Anti-miR-128 promotes mitochondrial gene and protein expression and represses an inflammatory transcriptional program in *mdx*^5c^*^v^*mouse muscle. (A) Functional category enrichment in upregulated genes (light gray) and down-regulated genes (dark gray) by anti-miR-128 treatment compared with scramble-treated *mdx*^5c^*^v^* mouse gastrocnemius muscle from cohort 1. Bars represent normalized enrichment score (NES) according to Gene Set Enrichment Analysis (GSEA). (H, Hallmark; R, Reactome; FDR<0.05) (B-D) GSEA enrichment plots of the Oxidative Phosphorylation Hallmark gene set (B), the TNFα Signaling via NFκB Hallmark gene set (C), and the Epithelial Mesenchymal Transition Hallmark gene set (D). (E) RT-qPCR showing upregulation of key genes for mitochondrial biogenesis, oxidative phosphorylation, fatty acid oxidation, energy sensors, and mitochondrial dynamics in gastrocnemius muscle from cohort 1. (F) RT-qPCR for *Tnfa* in gastrocnemius muscle from cohort 1. (G) Western blot for PGC-1α, PPARα, CPT1b, and β-tubulin in soleus muscle from mdx^5c^^v^ mice treated with scramble or anti-miR-128 from cohort 1 and quantified in (H). Data are presented as mean +/- SD. N= 5 per group (A-E); N=4-5 per group (F); N=4 per group (G-H). ns, not significant, *p<0.05, **p<0.01, ***p<0.001 (unpaired, two-tailed, t-test (E, F, H))

### Anti-miR-128 Improves Cardiac Function and Myocardial Proteome in a Porcine Model of DMD

DMD patients suffer from dilated cardiomyopathy, and with improvements in respiratory care, heart failure is now the leading cause of death.^41–44^ As *mdx*^5c^*^v^*mice do not fully recapitulate DMD-associated cardiac dysfunction, we next evaluated translational relevance of miR-128-3p inhibition on cardiac dysfunction in the *DMD^Y/−^* pig model.^45^ Six-week-old pigs were treated for 8 weeks with anti-miR-128 or scramble at 5 mg/kg (Fig. 6A). One animal in the anti-miR-128 treatment group died of unknown cause prior to study completion and was not included in endpoint analyses. Anti-miR-128 treatment efficiently suppressed miR-128-3p in myocardium of *DMD^Y/-^* pigs (Fig. 6B). Scramble-treated *DMD^Y/-^* pigs exhibited an increase in serum troponin levels due to cardiomyocyte damage (Fig. 6C), while those treated with anti-miR-128 showed a trending decrease, consistent with improved cardiomyocyte integrity. Cardiac function was measured by echocardiography at baseline and after 8 weeks of treatment. Given the phenotypic heterogeneity of the *DMD^Y/-^* model, we assessed functional trajectories at the individual animal level. Despite variable baseline cardiac function, all anti-miR-128-treated animals showed improvement from baseline over the treatment period, whereas scramble-treated animals showed a mixed or declining trajectory (Fig. 6 D-E). Using a linear mixed-effect model to compare anti-miR-128- and scramble-treated groups, the group-by-time interaction is significant for both EF and FS, indicating that the anti-miR-128 animals improved significantly more over the treatment period than the scramble control animals. Cardiac function was higher in the scramble-treated group at baseline, but by the end of the treatment period, this difference was no longer significant, while anti-miR-128-treated animals had improved significantly from their baseline. Serum chemistry shows increases in serum ALT and LDH of *DMD^Y/-^*pigs that is consistent with the underlying disease phenotype (Table 2). While no broad pattern of treatment-related abnormalities was identified, the small sample size precludes a definitive safety conclusion. Together, these data demonstrate that anti-miR-128 preserves cardiac function in a large animal model of DMD cardiomyopathy and support its translational potential.

**Figure 6.**
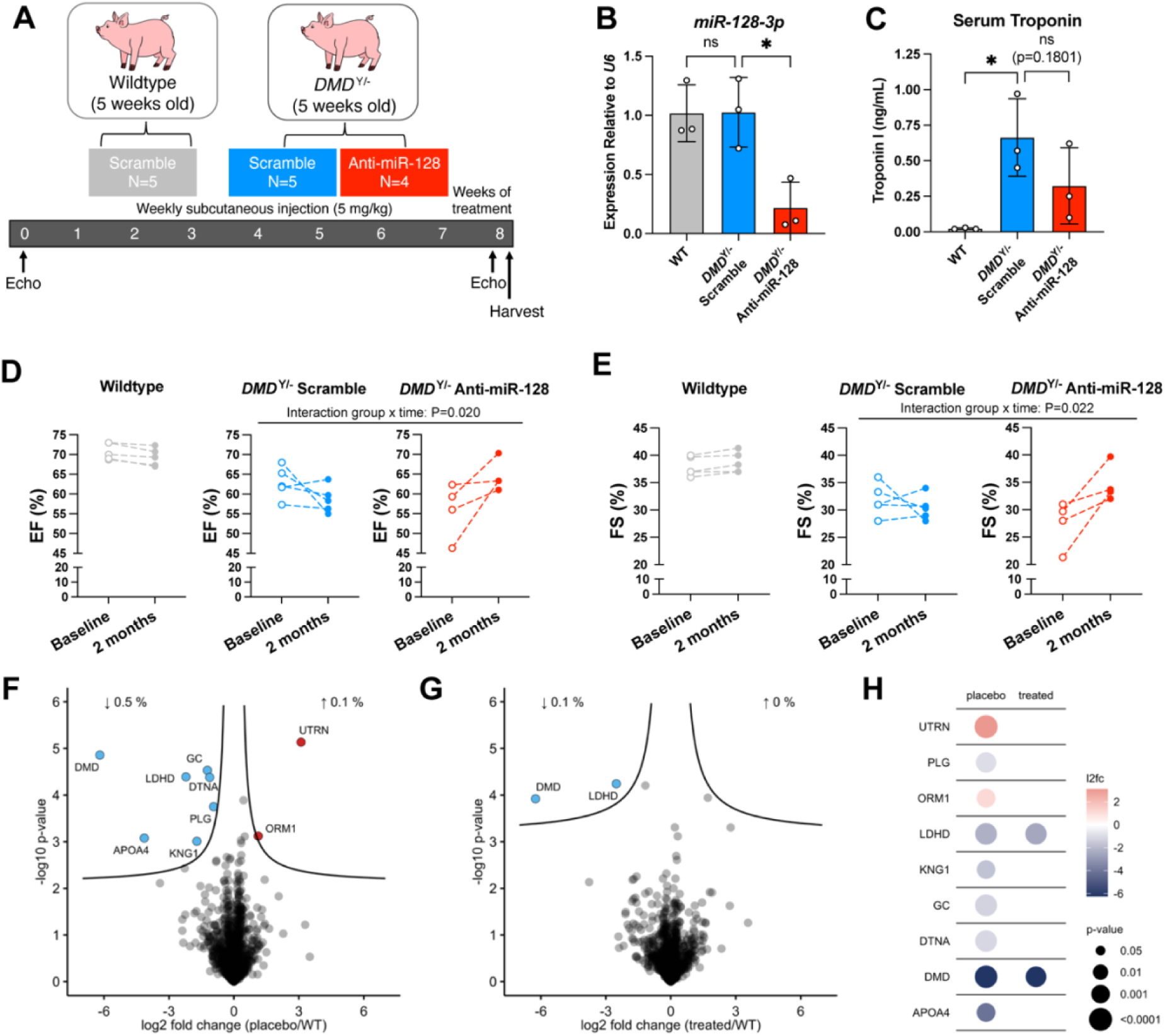
Anti-miR-128 improves cardiac function and myocardial proteome of *DMD^Y/-^* pigs. (A) Schematic of experimental design. *DMD^Y/-^* pigs were treated from 6 weeks of age with anti-miR-128 or scramble at 5 mg/kg for 8 weeks. Echocardiography was performed at baseline (5 weeks of age) and 8 weeks of treatment to assess cardiac function. (B) *miR-128-3p* expression relative to *U6* from myocardium of wildtype and *DMD^Y/-^* pigs treated with anti-miR-128 or Scramble. (C) Serum cardiac troponin I (cTnI) levels at 8 weeks of treatment. (D) Ejection fraction (EF) of wildtype, *DMD^Y/-^* scramble-treated, and *DMD^Y/-^* anti-miR-128-treated pigs at baseline and 8 weeks of treatment. (E) Fractional shortening (FS) of wildtype, *DMD^Y/-^* scramble-treated, and *DMD^Y/-^* anti-miR-128-treated pigs at baseline and 8 weeks of treatment. (F) Volcano plot visualization of myocardium proteome alterations. Proteins significantly altered in abundance in scramble treated *DMD^Y/-^* pigs compared to wildtype (FDR<0.05) are colored in blue for downregulation and red for upregulation. (G) Volcano plot visualization of myocardium proteome alterations. Proteins significantly altered in abundance in anti-miR-128-treated *DMD^Y/-^* pigs compared to wildtype pigs (FDR<0.05) are colored in blue for downregulation and red for upregulation. (H) Dot plot visualization of myocardium proteome alterations. Proteins significantly altered in abundance in scramble-treated *DMD^Y/-^* pigs compared to wildtype (left) and anti-miR-128-treated *DMD^Y/-^* pigs compared to wildtype (right) are colored in blue for downregulation and red for upregulation. Data are presented as mean +/-SD (B-C) and values for each animal at baseline and 8 weeks (D-E). N=3 per group (B-C) and N=4-5 per group (D-H). *p<0.05, **p<0.01, ***p<0.001 (one way ANOVA followed by Dunnett’s multiple-comparisons test (B-C), linear mixed-effects models with group, time, and group x time as fixed effects and animal as random effect, df=7 in both models (D-E)).

To gain insight into the molecular changes accompanying improved cardiac function in DMD pigs, we performed proteomics analysis of the myocardium. Proteomics of the *DMD^Y/-^* compared with wildtype myocardium confirmed the expected reduction in DMD protein. Other downregulated proteins in scramble-treated *DMD^Y/-^* animals included those involved in lipid transport and metabolism (APOA4), wound healing (PLG, KNG1), dystrophin associated complex proteins (DTNA), mitochondrial lactate metabolism (LDHD), and vitamin D binding protein (GC) (Fig. 6F, H). Increased proteins included utrophin (UTRN) and inflammation marker (ORM1) (Fig. 6F, H). When examining the proteome of anti-miR-128-treated *DMD^Y/-^* versus wildtype animals, most of these aberrantly expressed proteins are no longer differentially expressed, though LDHD, and DMD remain changed (Fig. 6G, H). This pattern is consistent with partial normalization of the disease-associated myocardial proteome following anti-miR-128 treatment. These data show that the proteomic signature associated with the disease state of the DMD pig model is ameliorated by anti-miR-128 treatment despite persistent dystrophin deficiency, extending the functional and molecular observations from *mdx*^5c^*^v^*mice to a large-animal model of DMD cardiomyopathy.

### Anti-miR-128 Reverses Aging-Associated Transcriptional Programs

Prior cross-species transcriptomic analyses of aging tissues have identified a conserved aging signature that is characterized by downregulation of mitochondrial function pathways and upregulation of inflammatory and immune pathways.^11^ This signature is reversed by longevity interventions such as caloric restriction, growth hormone deficiency, and rapamycin. We asked whether the convergent changes in transcriptional programs observed with anti-miR-128 treatment across sarcopenia, MI, and DMD models resemble this anti-aging program. Strikingly, the transcriptional profiles of anti-miR-128-treated aged mouse gastrocnemius muscle, post-I/R MI mouse heart, and *mdx*^5c^*^v^* gastrocnemius muscle closely resembled the gene expression profiles of long-lived animals and humans as well as longevity interventions, as shown by pathway enrichment scores across conditions (Fig. 7, S6). These data suggest that anti-miR-128-treated muscle tissues exhibit an anti-aging transcriptional signature, providing a unifying mechanistic framework for the functional benefits observed across the distinct striated muscle diseases studied here.

**Figure 7.**
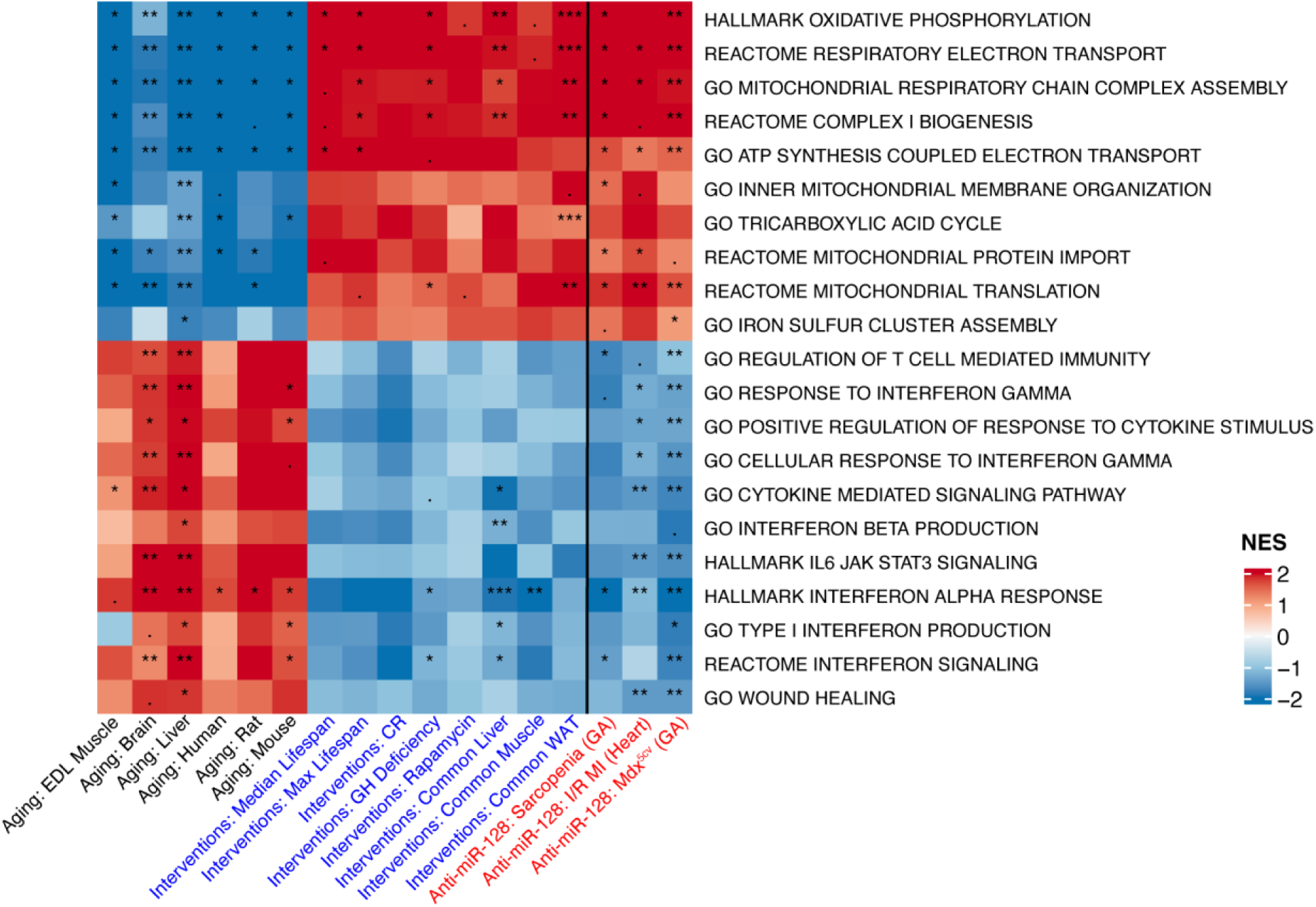
Anti-miR-128 reverses the transcriptional hallmarks of muscle aging and recapitulates longevity intervention signatures across disease contexts. Comparison of gene set enrichment analysis of the expression profile of aged C57BL/6J gastrocnemius muscle, *mdx*^5c^*^v^* gastrocnemius muscle, and post-I/R cardiac tissue treated with anti-miR-128 (red text) compared to published translational signatures of mammalian aging (black text) and signatures of longevity promoting interventions (blue text). For each experimental model, enrichment was calculated relative to the corresponding scramble-treated control. Labels in red denote the anti-miR-128 treatment datasets, labels in black denote published mammalian aging signatures, and labels in blue denote published longevity-intervention signatures. Red boxes indicate upregulated pathways, and blue boxes indicate downregulated pathways within each dataset. ***P.adjusted < 0.001; **P.adjusted < 0.01; *P.adjusted < 0.05; . P.adjusted < 0.1

## Discussion

In this study, we identify miR-128-3p as a regulator of a conserved aging-associated program shared across aging-related sarcopenia, myocardial infarction, and Duchenne muscular dystrophy. Despite substantial differences in etiology, all three conditions exhibited a common transcriptional response to miR-128-3p inhibition, characterized by activation of mitochondrial metabolism and suppression of inflammatory, fibrotic, and mesenchymal gene programs. Functional improvement accompanied these molecular changes in every disease context, indicating that diverse forms of striated muscle dysfunction, including physiological aging, acute ischemic injury, and inherited muscle degeneration, converge on a common tissue state.

Mitochondrial dysfunction and inflammation are tightly coupled processes that reinforce one another. Damaged mitochondria generate reactive oxygen species, release mitochondrial DNA into the cytosol, and activate innate immune sensors including cGAS-STING, TLR9, and the NLRP3 inflammasome.^46–51^ Inflammatory signaling, in turn, further impairs mitochondrial dynamics, oxidative phosphorylation, and redox homeostasis, generating a feed-forward loop that sustains tissue dysfunction.^9^ Recent studies have shown that mtDNA driven activation of cGAS-STING promotes chronic inflammation and a type I interferon signature during aging.^52^ Our data are consistent with a model in which miR-128-3p acts upstream of this loop: miR-128-3p inhibition induces genes governing mitochondrial biogenesis, fatty acid oxidation, mitochondrial dynamics, mitophagy, and lysosomal function, while consistently suppressing inflammatory, fibrotic and EMT programs. This control is exerted at multiple nodes of metabolism, including insulin signaling through INSR and IRS1, longevity signaling through SIRT1, mitochondrial biogenesis through PGC1α and TFAM, fatty acid oxidation through PPARα, mitochondrial dynamics through OPA1, and mitophagy and lysosomal function through ULK1, PINK1, and TFEB.^25,28,39,40^ These, together with the complex I subunit NDUFS4, have been validated as direct miR-128-3p targets.^25,28,39,40^ Together, these observations indicate that miR-128-3p coordinates multiple integrated metabolic and mitochondrial quality control programs rather than isolated pathways.

The transcriptional changes induced by anti-miR-128 resemble those of established longevity interventions. Multi-species transcriptomic analyses have shown that aging is accompanied by repression of oxidative phosphorylation and TCA cycle activity together with activation of inflammatory pathways, particularly interferon-α and interferon-γ signaling.^10–12^ These age-associated signatures are reversed by caloric restriction, rapamycin, growth hormone deficiency, and heterochronic parabiosis. ^10–12^ Across aged skeletal muscle, infarcted myocardium, and dystrophic muscle, anti-miR-128 elicited a similar transcriptional shift characterized by enhanced mitochondrial metabolism and attenuated inflammatory signaling, with interferon responses prominent among the suppressed pathways. These findings suggest that miR-128-3p participates in a conserved aging-associated molecular program and that the functional benefits of its inhibition across diverse disease contexts may arise, at least in part, from modulation of this program. From this perspective, myocardial infarction and Duchenne muscular dystrophy may be viewed, in part, as pathological states that accelerate molecular features of tissue aging.

Within this framework, aging-related sarcopenia, myocardial infarction, and Duchenne muscular dystrophy can be regarded as distinct manifestations of muscle stress that differ principally in acuity and chronicity. At the chronic extreme, the accumulation of mild damage that is no longer efficiently repaired produces progressive mitochondrial decline and chronic inflammatory activation in aging muscle.^53–55^ A recent analysis of human tissue supports the relevance of this program to muscle aging in our own species. Using histology-derived clocks of biological age across 40 tissues in the Genotype-Tissue Expression cohort, and gene expression from the same donors, accelerated biological aging of human skeletal muscle was associated with reduced expression of oxidative phosphorylation, fatty acid metabolism and peroxisomal gene sets and with elevated interferon alpha and gamma, epithelial-mesenchymal transition, transforming growth factor beta and complement programs.^53^ Each of these eight gene sets moves in the direction opposite to the anti-miR-128 response reported here. The ability of anti-miR-128 to improve function across all three models indicates that this downstream state is therapeutically tractable even when the initiating insult cannot be removed. This principle is illustrated clinically by corticosteroids in Duchenne muscular dystrophy, which slow functional decline by suppressing inflammation without correcting the underlying dystrophin deficiency.^56,57^

Beyond mitochondrial dysfunction and inflammation, our data suggest that miR-128-3p may also regulate a broader process involving erosion of cellular identity. Loss of PRC2 function, epigenetic drift, and declining lineage fidelity are increasingly recognized features of aging tissues and have been proposed to underlie mesenchymal drift, a convergent aging process in which cells progressively acquire fibrogenic characteristics at the expense of specialized identity.^54–59^ Consistent with this framework, epithelial-mesenchymal transition, extracellular matrix organization, and TGFβ signaling gene sets were among the most consistently downregulated signatures across all three RNA-seq datasets, while oxidative phosphorylation was significantly upregulated following anti-miR-128 treatment. Fibrosis was also reduced histologically in both the myocardial infarction and dystrophic muscle models, suggesting that miR-128-3p inhibition may partially limit the shift toward a mesenchymal, fibrogenic state. In the context of myocardial infarction, miR-128-3p has been shown to directly target SUZ12, a core subunit of PRC2, linking this microRNA to chromatin-level epigenomic control of cellular state. ^27^ Although we did not detect induction of the *Suz12* transcript itself, anti-miR-128 treatment was associated with coordinated changes in SUZ12/Polycomb-linked transcriptional programs in the post-MI heart. These findings are consistent with post-transcriptional de-repression of SUZ12, potentially enhancing PRC2-mediated repression of downstream target genes. One possibility is translational de-repression of SUZ12 without a change in transcript level, although SUZ12 protein was not measured here. However, these observations raise the possibility that the degenerative state regulated by miR-128-3p in striated muscle represents a tissue- or cell type-specific manifestation of mesenchymal drift, and that its inhibition favors a more metabolically active and identity-preserved cellular state. ^60,61^

The evolutionary history of miR-128-3p offers a framework for understanding why such a program might be conserved. The miR-128-1 locus sits within a positively selected haplotype on chromosome 2q21.3 associated with energy storage and metabolic efficiency across multiple mammalian species. ^25^ In environments marked by periodic nutrient scarcity, a program that promotes energy conservation and restrains energy expenditure could confer a survival advantage. However, when persistently engaged during aging or chronic tissue injury, the same adaptive program may become maladaptive, contributing to mitochondrial dysfunction and chronic inflammation observed across aging and striated muscle diseases. This interpretation is consistent with the concept of antagonistic pleiotropy, whereby traits that enhance fitness earlier in life contribute to late-life decline, and miR-128-3p regulates one such program in striated muscle.

Several features of these results support translational potential. Anti-miR-128 was tolerated with chronic dosing in rodents, pigs, and non-human primates and improved muscle function across aging-related sarcopenia, acute cardiac injury, and genetic muscular dystrophy, including in a large-animal model that better recapitulates human cardiac physiology than rodents. Human genetic associations of the miR-128 loci with grip strength and pulmonary function further support the relevance of this pathway to human muscle health. Notably, anti-miR-128 administered during the early period after ischemic injury improved cardiac function following MI, highlighting potential clinical therapeutic relevance in acute coronary syndromes, where prevention of adverse remodeling could reduce progression to heart failure, which affects 14-36% of MI patients.^62,63^ Systemic administration of naked ASO in saline was sufficient to target both skeletal and cardiac muscle without conjugation or uptake-enhancing modifications; emerging muscle-targeting strategies, including transferrin receptor 1-directed antibodies and peptides, may enable tissue-specific dissection of the relative contributions of individual organs to these effects. ^64–66^ Because we and others have established roles for miR-128-3p in liver and adipose tissue as well, such approaches will also help determine whether the functional improvements reported here are predominantly muscle-intrinsic or arise in part through crosstalk among metabolic tissues. Together, these findings position miR-128-3p inhibition as a means of shifting striated muscle out of the tissue state common to these diseases, through a single post-transcriptional regulator.

### Limitations of Study

While our findings support the therapeutic potential of anti-miR-128 for modulating an aging-associated program shared across multiple diseases of striated muscle, several limitations and questions remain. We did not assess lifespan following chronic anti-miR-128 administration, and it therefore remains unknown whether reversal of the transcriptional aging signature translates into extension of lifespan in addition to healthspan. The mechanistic relationship between mitochondrial dysfunction and inflammation also remains to be defined, including whether miR-128-3p directly regulates inflammatory pathways or whether its effects on inflammation are secondary to mitochondrial restoration. Furthermore, single-cell and single-nucleus analyses, together with muscle-specific delivery approaches, will be important for determining whether the observed benefits arise primarily from muscle-cell-intrinsic effects or involve immune and other non-muscle cell populations. These approaches may also clarify the contribution of metabolic tissues, including liver and adipose tissue, to the systemic effects of miR-128-3p inhibition on striated muscle function.

Our studies were constrained by the available disease models and the feasibility of addressing all questions within a single study. The *mdx*^5c^*^v^*mouse model has a milder phenotype than human DMD, and treatment was initiated at 3 weeks of age, prior to overt pathology, so these findings reflect a preventative rather than therapeutic paradigm. Complementary studies in the *DMD^Y/-^* model, in which treatment began after cardiac dysfunction was established, provided evidence that anti-miR-128 also improves cardiomyopathy, a major feature of the human disease. The *DMD^Y/-^* model, however, exhibits substantial phenotypic variation, underscoring the importance of validating these findings in additional models. As DMD is X-linked and predominantly affects males, these studies were conducted in male animals. Similarly, although we used well-established surgical models of MI, these experiments were performed in young male mice, whereas MI occurs predominantly in older individuals of both sexes. Given that anti-miR-128 improves post-MI cardiac function in young mice while reversing an aging-associated transcriptional program, it will be important to determine whether its effects are enhanced in aged animals in which this program is already engaged before ischemic injury. Finally, our aging-related sarcopenia studies were performed exclusively in male mice because of the cost and extended duration required for studies in aged mice. We prioritized sufficient sample sizes to maintain statistical power for the primary endpoints, but future studies should incorporate female mice, particularly given established sex-specific differences in muscle aging and adaptation.

Our non-human primate tolerability study was likewise limited to small, male only groups as well as group allocation by NASH score and liver lipid content rather than randomization leaving analytes imbalanced between arms. Complement activation products and coagulation parameters as well as higher doses and longer exposure are needed to further assess ASO tolerability to support clinical development.

## Author contributions

Conceptualization, A.M.N., M.L.S., M.A.B., L.X., A.W., and S.Kau.; Methodology, M.A.B., L.X., X.W., M.S., M.B.L., A.P., P.I.H., A.L., B.S., T.F., D.O.R., J.C.W.H., G.B., L.C., T.S., and R.I.S.; Investigation, M.A.B., L.X., X.W., M.S., C.Z., J.Y.L., R.S., S.D.V., M.B.L., A.P., C.G., F.G., L.C., P.I.H., M.V., R.A., C.P., C.J., D.O.R., J.C.W.H., A.L., B.S., T.F., and E.W.; Formal Analysis, M.A.B., L.X., X.W., G.B., C.J., T.M.S., D.O.R., J.C.W.H., B.S., T.F., and R.I.S.; Data Curation, G.B., R.I.S., D.O.R., J.C.W.H., B.S., and T.F.; Resources, N.K., E.W., R.J.P., A.F., R.E.T., A.P., S.Kau., S.Kan., and M.L.S.; Validation, M.A.B., L.X., X.W., and M.S.; Visualization, M.A.B., L.X., X.W., G.B., C.J., and B.S.; Writing – Original Draft, A.M.N., M.A.B., and L.X.; Writing – Review & Editing, all authors; Supervision, A.M.N., M.L.S., E.W., T.F., R.J.P., R.E.T., R.I.S., S.Kan., S.Kau., and J.W.K.; Project Administration, A.M.N., L.X., A.F., and E.W.; Funding Acquisition, A.M.N., E.W., S.Kau., R.E.T., S.Kan., and M.L.S.

## Declaration of Generative AI and AI-Assisted Technologies in the Writing Process

During the preparation of this work, the authors used the AI tools Claude (Anthropic) and ChatGPT (OpenAI) to review the manuscript and suggest edits. After using these tools, the authors reviewed suggestions and edited content as needed and take full responsibility for the content of the publication. Generative AI was not used to create or alter any figure, image, or graphical abstract and was not used to generate or analyze research data.

## Acknowledgements

We thank Denise Schichnes and Steven Ruzin at The RCNR Biological Imaging Facility at the University of California, Berkeley for their microscopy technical support; Reena Zalpuri and Danielle Jorgens at the Electron Microscope Lab at UC Berkeley for their electron microscopy support; and Kevin Siao at the UCSF Liver Center for histology services. These studies were supported by the following grants: National Institutes of Health/NIDDK R01 DK114277 to A.M.N.; NIH/NINDS R56 NS128721 to A.M.N. and S.Kan.; a Novo Nordisk Foundation Challenge Grant (NNF18OC0033438) to S.Kau., A.M.N., and R.E.T.; NIH/NIDDK DK147451 and DK116008 to S.Kan; NIH/NIDDK P30 DK040561 to R.I.S.; Wellcome Trust (101550/Z/13/Z) to R.J.P; J.W.K. was supported by the NIH: P30 DK116074 (to the Stanford Diabetes Research Center), R01 DK116750, R01 DK120565, R01 DK106236, and R01 DK137889. T.M.S. was supported by a grant from the Novo Nordisk Foundation (NNF19OC0054265) and the Stanford Bio-X program. We thank the Barrett, Hathaway, Killian, and Westly families for generous support. E.W. was supported by grants from the German Center for Child and Adolescent Health (DZKJ; 01GL2406A), Else Kröner-Fresenius Foundation (2015_180, 2018_T20, 2022_EKSE.41), and Leducq Foundation (23CVD01). M.L.S. was supported by NIH grant U54 HL147127. A.M.N. and M.L.S. were also jointly supported by a grant from the UC Berkeley Bakar Fellows Program (Spark Award).

This work is dedicated to the memory of Dr. Matthew L. Springer, whose scientific insight and collaboration were critical to this study.

## Methods

### Key Resources Table

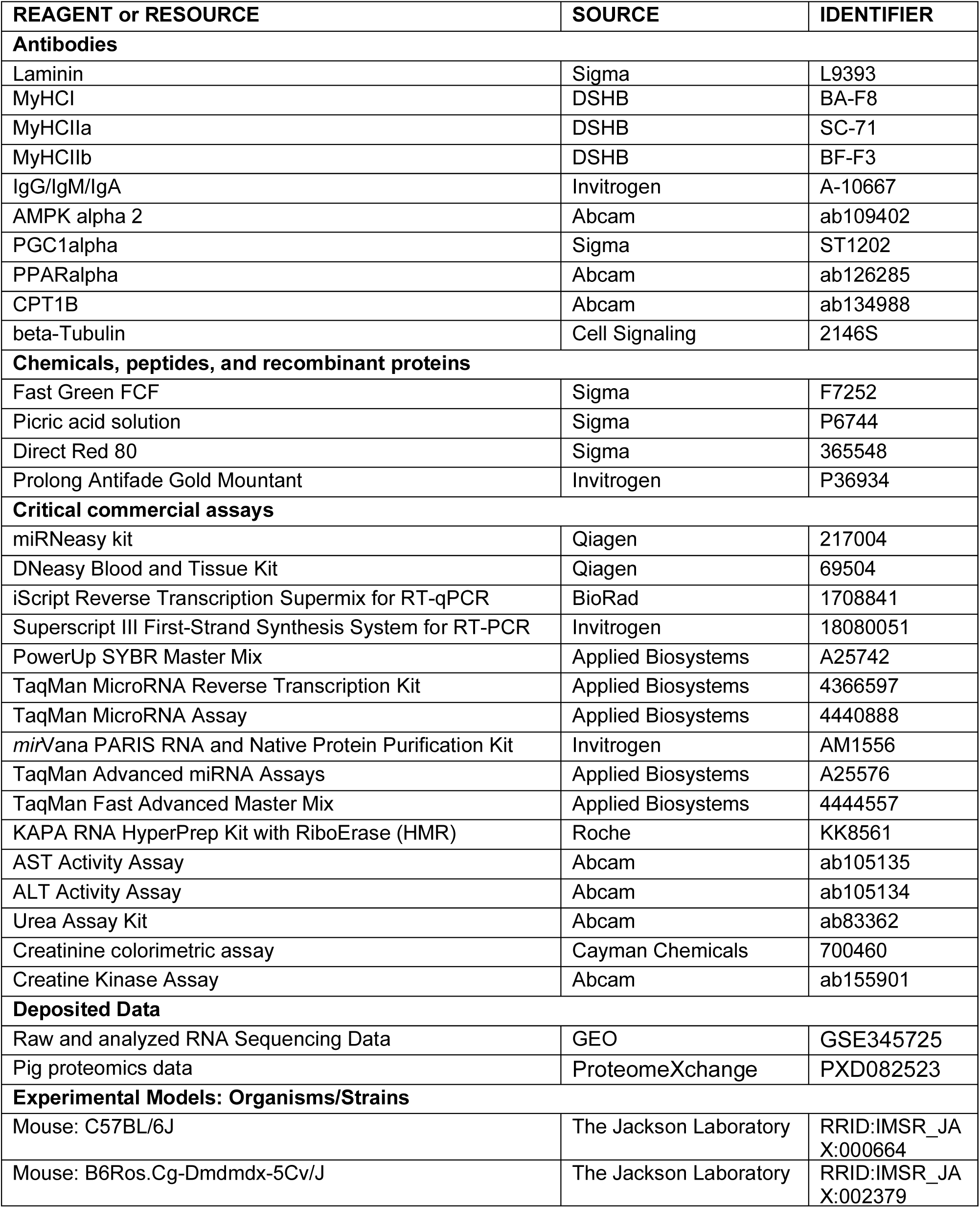
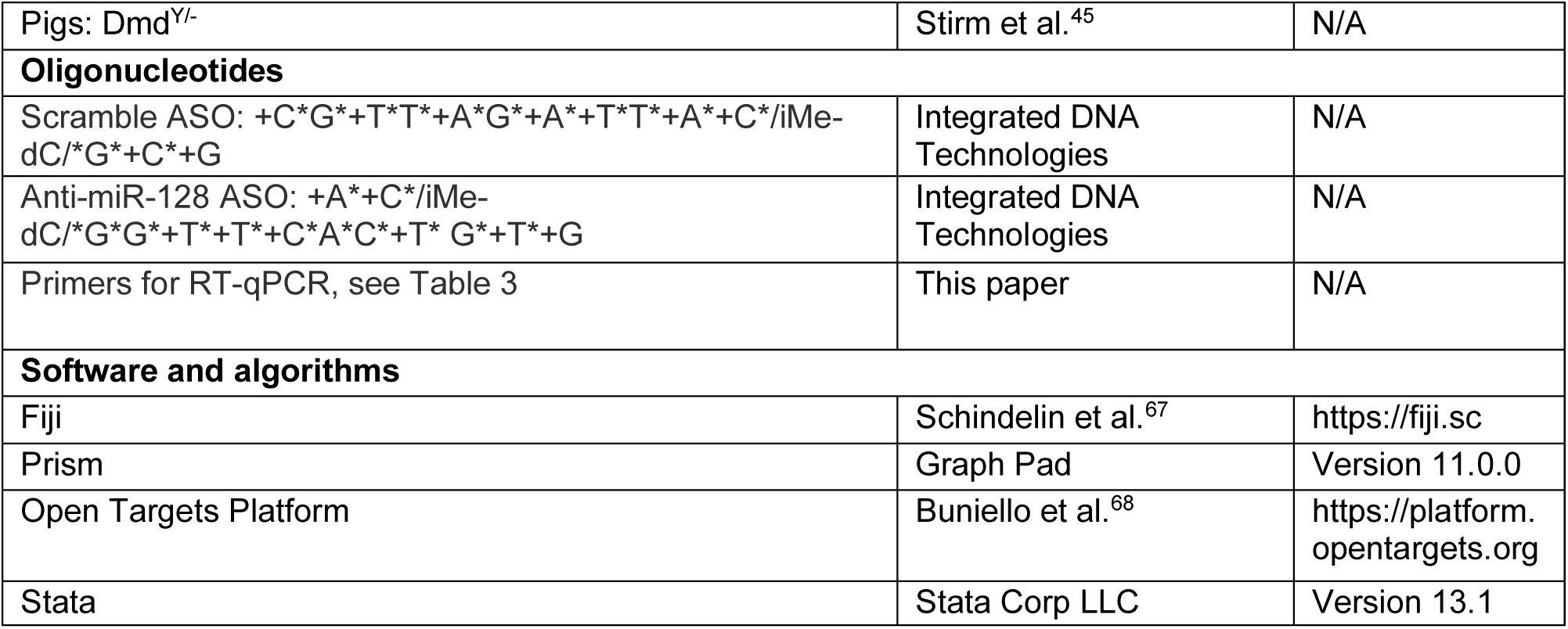

## Resource Availability

### Lead contact

Further information and requests for reagents may be directed to, and will be fulfilled by, the lead contact, Anders M Näär.

### Materials availability

All materials are commercially available and can be purchased directly from vendors.

### Data and code availability

- RNA sequencing data have been deposited in the Gene Expression Omnibus and are publicly available as of the date of publication. The three datasets (aged gastrocnemius, post-I/R left ventricle, *mdx*^5c^*^v^*gastrocnemius) are linked under accession GSE345725.
- Proteomics data are available via ProteomeXchange with identifier PXD082523.
- GWAS credible set data are available from the Open Targets Platform, and the trait label map are included in supplementary documents (Table 4).
- Any additional information required to reanalyze the data reported in this paper is available from the lead contact upon request.

## Experimental Model and Subject Details

### Mice

C57BL/6J mice (strain #000664) were purchased from the Jackson Laboratory, Bar Harbor, ME. Male mice for aging studies were aged 85 weeks and those for myocardial infarction studies were aged 9 weeks. *Mdx*^5c^*^v^* mice (Strain #002379) and wildtype C57BL/6J controls (strain #000664) were obtained from the Jackson Laboratory and breeding was conducted at UC Berkeley’s Northwest Animal Facility. All experimental procedures were conducted in accordance with IACUC regulation and were approved by UC Berkeley (AUP-2018-10-11513-2) or UC San Francisco’s IACUC.

### Pigs

The porcine DMD model, with a deletion of *DMD* exon 52, has been described in detail elsewhere.^45^ All pig experiments were performed according to the German Animal Welfare Act and Directive 2010/63/EU on the protection of animals used for scientific purposes and were approved by the responsible animal welfare authority (Government of Upper Bavaria; permission 55.2-1-54-2532-163-2014). All pigs were kept in a specified pathogen-free environment (Center for Innovative Medical Models) at the Ludwig-Maximilian University Munich. Food and water were provided ad libitum.

### Cynomolgus Monkeys

Adult male cynomolgus monkeys were used for non-human primate studies at the University of Kentucky. All experiments were performed according to University of Kentucky’s IACUC regulations and approved by their IACUC.

### Dogs

Dogs (*Canis familiaris*) were from the DE50-MD colony, housed in a dedicated facility at the Royal Veterinary College (RVC), London. ARRIVE guidelines were followed at all times. The study was conducted within project license P9A1D1D6E (granted 11 June 2019) assigned under the Animal (Scientific Procedures) Act 1986, according to UK legislation and approved by the Royal Veterinary College local Animal Welfare Ethical Review Body.

### DMD Patients

Serum samples were obtained from patients seen in the pediatric neuromuscular clinic at UCSF Benioff Children’s Hospital under IRB protocol 22-36067. Informed consent was obtained from at least one parent or legal guardian and assent from patients 7 years of age and above, if able. The analyzed cohort samples include 8 boys with Duchenne muscular dystrophy (all male; mean age 10.5 years, range 2–17), and 2 comparison participants (2 male; ages 3 and 10 with cerebral palsy diagnosis). Of the DMD participants, 4 were ambulatory and 4 non-ambulatory; 6 were receiving corticosteroids (5 prednisone or prednisolone, 1 deflazacort), 1 was receiving exon-skipping therapy, and 2 were receiving neither. Comparison participants were patients of the same clinic diagnosed with cerebral palsy; they are therefore non-muscle disease controls rather than healthy controls. Blood was collected into the same tube type, processed by the same procedure, and stored for comparable durations for all participants under the single IRB protocol.

## Method Details

### Locked Nucleic Acid Antisense Oligonucleotides

Locked nucleic acid modified antisense oligonucleotides (LNA ASOs) targeting miR-128-3p (+A*+C*/iMe-dC/*G*G*+T*+T*+C*A*C*+T* G*+T*+G) or scramble control (+C*G*+T*T*+A*G*+A*+T*T*+A*+C*/iMe-dC/*G*+C*+G) were purchased from Integrated DNA Technologies and used for all anti-miR-128 animal experiments. (+ denotes Affinity Plus (locked nucleic acid base); * denotes Phosphorothioate bonds; and /iMe-dC/ denotes 5-Methylcytosine (5-methyl-dC)). LNA ASOs were delivered dissolved in saline and administered by subcutaneous injection.

### Mouse Sarcopenia Studies

85-week-old male C57BL/6J mice (strain #000664) were procured from the Jackson Laboratory, Bar Harbor, ME. Mice were acclimated in UC Berkeley’s Northwest Animal Facility for one week prior to commencing treatment. Anti-miR-128-3p antisense oligonucleotide (ASO) or scramble control ASO was dissolved in saline and delivered by subcutaneous injection at 10 mg/kg once per week. Treatment duration was 20 weeks for cohort 1 and 16 weeks for cohort 2. Muscle function was measured longitudinally throughout the duration of the study to monitor functional decline and the therapeutic effect of anti-miR-128. Body composition was measured at 15 weeks of treatment for cohort 2. Upon sacrifice, tibialis anterior, quadriceps, and gastrocnemius muscles were collected and about 1 mL of blood was collected by cardiac puncture. Tissues were snap frozen in liquid nitrogen, prepared for histology as described below, or embedded in OCT for sectioning. Blood was centrifuged for 5 minutes at 4°C to obtain serum. Serum and cryopreserved tissues were stored at -80°C until analysis. All mouse procedures were approved by the University of California, Berkeley Institutional Animal Care and Use Committee. All protocols conform to federal regulations, the National Research Council Guide for the Care and Use of Laboratory Animals, and the Public Health Service Policy on Humane Care and Use of Laboratory Animals.

### Surgical Mouse Myocardial Infarction Studies

Male C57BL/6J mice (10 weeks old) were obtained from The Jackson Laboratory (Bar Harbor, ME). For the ischemia/reperfusion (I/R) myocardial infarction (MI) model, as previously described,^69^ the mice were anesthetized with 2% isoflurane and received buprenorphine (0.1 mg/kg, subcutaneous) before surgery, after surgery, and once daily for 3 days for postoperative analgesia. Mice were intubated and mechanically ventilated using a volume-controlled small animal ventilator (Harvard Rodent Ventilator, Model 683, South Natick, MA). Following thoracotomy, the heart was exposed and the left anterior descending (LAD) coronary artery was occluded for 30 minutes to induce regional myocardial ischemia. Briefly, a 7-0 nonabsorbable suture was passed under the LAD approximately 2 mm from the tip of the left auricle and tied to achieve temporary occlusion. After 30 minutes, the ligation was released to allow reperfusion of the ischemic myocardium. The chest was closed after removal of residual air from the thoracic cavity. Mice were monitored every 15 minutes during recovery until fully ambulatory. For the permanent LAD ligation MI model, as previously described^70^ mice underwent the same surgical procedure except that the LAD was permanently ligated. Procedural mortality averaged 3% for the I/R MI model and 10% for the permanent ligation MI model.

### Canine Duchenne Muscular Dystrophy Studies

Dogs (*Canis familiaris*) were from the DE50-MD colony, housed in a dedicated facility at the Royal Veterinary College (RVC), London. Wild type (WT) and DE50-MD dogs were housed typically in groups of 2–3 (groups determined according to dog temperament and hierarchies). Dogs were housed in indoor kennels (4.5 m^2^ to 7.5 m^2^) according to the Animal (Scientific Procedures) Act 1986 of the United Kingdom (ASPA) code of practice (Section 2) with 12-hour light/dark cycle, 15–24°C) and given daily access to large outdoor paddocks (approximately 100 m^2^). Dogs were produced by breeding carrier female Beagles (RCC strain, Marshall Bioresources)-cross dogs, with WT male Beagles (RCC strain). Dogs were observed daily by animal technician staff, and any concerns were reported to the Study director, Named Veterinary Surgeon and Named Animal Care and Welfare Officer. All efforts were made to minimize any animal suffering throughout the study. Humane endpoints were established before commencing the study: dehydration (unresolved by fluid treatment), lethargy/motor dysfunction, weight loss/dysphagia, dyspnea, listless behavior/demeanor or heart failure. None of the dogs included in this study reached any humane endpoint during the 18-month study period. At seven days of age, puppies were microchipped and cheek swabs were taken for preparation of DNA (GeneJET genomic DNA kit, #K0721, Thermofisher^TM^) for genotyping. Polymerase chain reaction (PCR) was performed on a Veriti 96 Well Thermal Cycler (Applied Biosystems) using primers spanning the DE50-MD mutation site (splice donor site of dystrophin exon 50): Forward primer 5’-3’ sequence AGCTCTGATTGGAAGGTGGT; Reverse primer 5’-3’ sequence ACCTCAGTGTTGTGCTTTTGA. PCR products were sent for Sanger sequencing (SupremeRun, Eurofins Scientific) using the forward primer only. Other than a genotype of WT male or DE50-MD male, no additional exclusion criteria or randomization strategies were used when recruiting dogs. Dogs that were not recruited to studies were rehomed. Staff and researchers involved in animal husbandry, data acquisition and data analysis were not blinded to genotype.

### Mouse Duchenne Muscular Dystrophy Studies

*Dmd^mdx-5Cv^* mice (Strain #002379; referred to herein as *mdx*^5c^*^v^*) were purchased from the Jackson Laboratory, Bar Harbor, ME and bred with C57BL/6J for at least six generations. C57BL/6J mice (strain #000664) were obtained from the Jackson Laboratory, then bred in house, and aged matched mice were used as wildtype controls. Male *mdx*^5c^*^v^* mice and wildtype controls were treated weekly by subcutaneous injection of scramble control or anti-miR-128 at 10 mg/kg starting from three weeks of age. Mice were exercised twice per week during the study by treadmill running at 9 m/min for 30 minutes.^71,72^ Muscle function behavioral tests were performed at the beginning of the third week of treatment for cohort 1 and the beginning of the sixth week of treatment for cohort 2. Cohort 1 was harvested at three weeks of treatment and cohort 2 at six weeks of treatment. Upon sacrifice, tibialis anterior, quadriceps, and gastrocnemius muscles were collected and about 1 mL of blood was collected by cardiac puncture under deep anesthesia. Tissues were snap frozen in liquid nitrogen or embedded in OCT for sectioning. Blood was centrifuged for 5 minutes at 4°C to obtain serum. Serum and cryopreserved tissues were stored at -80°C until analysis. All mouse procedures were approved by the University of California, Berkeley Institutional Animal Care and Use Committee. All protocols conform to federal regulations, the National Research Council Guide for the Care and Use of Laboratory Animals, and the Public Health Service Policy on Humane Care and Use of Laboratory Animals.

### Pig Duchenne Muscular Dystrophy Studies

To standardize the experimental groups, all DMD piglets in the treatment and control groups as well as wildtype piglets were generated by breeding the same wildtype boar with heterozygous carrier sows (*DMD*^+/-^). One animal from the treatment group died for unknown reasons before the experiment was completed and could not be included in the measurements and sample collection. Baseline echocardiography was performed at five weeks of age (one week after weaning). DMD (male) pigs were randomly assigned to the treatment or control groups. Pigs were treated at six weeks of age with scramble control or anti-miR-128 at 5 mg/kg once per week for a duration of eight weeks. Injection was delivered subcutaneously into the skin fold just behind the auricle. Echocardiography was again performed at eight weeks of treatment. Blood samples were obtained at two weeks and eight weeks of treatment. Animals were euthanized, and necropsy and tissue sampling occurred after eight weeks of treatment. All pig experiments were performed according to the German Animal Welfare Act and Directive 2010/63/EU on the protection of animals used for scientific purposes and were approved by the responsible animal welfare authority (Government of Upper Bavaria; permission 55.2-1-54-2532-163-2014). All pigs were kept in a specified pathogen-free environment at the Center for Innovative Medical Models at the Ludwig-Maximilian University Munich.

### Mouse Dosing Studies

Eight-week-old C57BL/6J mice were given a single dose of anti-miR-128 and miR-128-3p levels in liver, gastrocnemius muscle, and cardiac muscle were measured 28 days later by RT-qPCR.

### Mouse Kinetics Studies

10-week-old C57BL/6J mice were given a single dose of anti-miR-128 by subcutaneous injection and serum and tissues were harvested 1, 4, 7, and 10 days later. For serum preparation, samples were collected at the indicated time points by tail tip incision and allowed to clot at room temperature for 1 hour. Samples were then centrifuged at 2,000 × g for 5 minutes at room temperature, and serum was collected from the supernatant. For tissue preparation, prior to euthanasia, mice underwent whole-body perfusion with 25 mL PBS. ASO levels were quantified by SplintR qPCR.

### Cynomolgus Monkey Toxicity Studies

Adult male cynomolgus monkeys (*Macaca fascicularis*) were used to evaluate the safety profile of chronic anti-miR-128 administration. Studies were conducted at the University of Kentucky under Institutional Animal Care and Use Committee (IACUC) oversight. Monkeys were pair-housed, when possible, in climate-controlled conditions with 12-h light–dark cycles. Animals were singly housed from ∼08:00-15:00 each day and at 08:00 and 11:00 received 75 g of a semi-synthetic fast-food diet (9% (by weight) Casein, 5% whey protein isolate 90, 15% sucrose, 20% fructose, 20% whole wheat flour, 16.4% lard, 0.2% fish oil, 0.19% cholesterol 6.31% cellulose, 2.5% Vitamin Mix, X75, 5% Mineral Mix, Hegsted IV, and 0.4% Calcium Carbonate), which provided on average 69 kcal/kg body weight/day. In place of water, a high fructose drink (60.5 g/L fructose, 49.5 g/L glucose, 10 mL/L tropical punch concentrate, and 0.2 mL/L red food coloring) was provided ad libitum to the animals. After 31 months of fast-food diet and high fructose drink consumption, monkeys were assigned into two groups based upon the MASH score and lipid content of liver biopsy samples collected at 24 months of fast-food diet feeding. Monkeys then received either anti-miR-128 (5 mg/kg) or vehicle (sterile saline) by subcutaneous injection (n=3 saline, 4 anti-miR-128 per group), administered once weekly for 24 weeks. Blood was collected from the femoral vein at baseline (week 0), weeks 1 and 2, and every 2 weeks thereafter through week 24. Prior to each blood collection, animals were fasted overnight, then anaesthetized with ketamine (5-10 mg/kg). At study termination (week 24), fasted monkeys were anaesthetized with ketamine (10 mg/kg), bled a final time, and euthanized by exsanguination with whole-body saline perfusion while anaesthetized with isoflurane (3–5% induction, 1–2% maintenance). Tissues were promptly removed and fixed in 10% neutral-buffered formalin for downstream histological analysis.

### GWAS Credible Set Analysis

To characterize the phenotypic landscape associated with the miR-128-1 and miR-128-2 loci, we retrieved genome-wide association study (GWAS) credible set data for the index variants rs1446585 (chr2:135,649,909, GRCh38; 2_135649909_A_G), rs1446584 (chr2:135,662,391, GRCh38; 2_135662391_A_G), rs1438307 (chr2:135,741,596, GRCh38; 2_135741596_G_T), and rs1513475 (chr3:35,682,764, GRCh38; 3_35682764_T_C) from the Open Targets Platform (https://platform.opentargets.org; accessed May 26, 2026).

For visualization, association p-values were converted to –log10(P-value) scale. Where multiple studies reported associations for the same trait or synonymous trait descriptions, only the most statistically significant association was retained per trait label. Associations were plotted as a PheWAS plot in which each point represents a unique labeled trait, ordered along the x-axis by statistical significance. Points were colored by broad trait category (lipids, anthropometric, lung function, physical performance, metabolic, cardiovascular event, or other). The genome-wide significance threshold (P < 5 × 10⁻⁸) is indicated by a dashed horizontal line. Trait labels were assigned from a manually curated mapping table (trait_label_map.csv) that groups synonymous GWAS trait descriptions into consolidated display labels. All analyses and visualizations were performed in R (version 4.4) using the dplyr, ggplot2, and ggrepel packages.

### Muscle Function Behavioral Tests

Muscle function was assessed using wire hanging or treadmill run time until exhaustion. For wire hanging, mice were acclimated to the testing room in their home cages then each mouse was gently placed with its forepaws hanging onto a 2 mm wire which is suspended between two supports over a cushioned supporting base. The time the mouse remained on the wire before falling was recorded. The test was repeated for a total of 3 trials, separated by 10-minute rest intervals, and performance was averaged across trials. Running distance to exhaustion was measured by either high-intensity or low-intensity running protocols using an Exer-6M treadmill (Columbus Instruments) as previously described.^73^ Mice were fasted for 2 hours prior to testing. For the low-intensity protocol, mice were trained to run for two days pre-test (day 1: 5 m/min for 10 min; day 2: 5 m/min for 1 min, 7.5 m/min for 1 min, 10 m/min for 8 min). On testing day, mice were run according to the following protocol: 5 m/min for 1 min; 8 m/min for 1 min; 12 m/min for 38 min; an additional 1 m/min faster every 10 min until 15 m/min; and +1 m/min every 5 min until 20 m/min is reached. For the high-intensity test, the running protocol was 5 m/min for 5 minutes then an increase of 1 m/min every minute until 20 m/min is reached. All running tests were conducted at a 0° incline. Exhaustion criteria were defined as any mouse that will not run for 20 seconds despite repeated gentle nudges.

### Gene and Protein Expression Analysis

Tissues were dissected, snap frozen in liquid nitrogen, and stored at -80°C until mRNA and protein extraction. For mRNA quantification, total RNA was extracted using the miRNAeasy kit (Qiagen) according to the manufacturer’s instructions. Reverse transcription to cDNA was performed using the Superscript III First-Strand Synthesis System for RT-PCR (Invitrogen) or iScript Reverse Transcription Supermix (BioRad). Gene expression was quantified by real-time PCR using PowerUP SYBR Green (Applied Biosystems) using the QuantStudio6 Real-Time PCR system (Applied Biosystems). mRNA CT values were normalized to r18S CT values measured in the same run. PCR primer sequences are provided in the Key Resources section. For protein expression, protein was extracted in RIPA buffer (50 mM Tris HCl pH 8.0, 150 mM NaCl, 1% NP-40, 0.5% sodium-deoxycholate, 0.1% SDS) with 1 mM DTT, 1 mM PMSF, and protease and phosphatase inhibitors (Pierce). Samples were homogenized, sonicated, and centrifuged for 10 min at 15,000 rpm and 4°C. Protein was quantified by BCA (Pierce). 30 µg of protein was prepared with Laemmli sample buffer (BioRad) and run on a 4-20% gradient Mini-Protean TGX Gel (BioRad). Samples were transferred to a PVDF membrane using the Transblot-Turbo Mini Transfer Pack system (BioRad). The membrane was blocked for 1-2 hours at room temperature in 5% milk in TBST. Primary incubation was overnight in 2.5% BSA in TBST with 0.02% sodium azide. The next day, the membrane was washed three times for 10 min in TBST. HRP-conjugated secondary antibody incubation was 2 hours at room temperature in 5% milk in TBST. Finally, the membrane was washed three times for 10 min in TBST. Membranes were visualized using Clarity Max Western ECL Substrate (BioRad) or SuperSignal West Pico PLUS Chemiluminescent Substrate (Pierce) with the Invitrogen iBright 1500 imaging system. Primary antibodies were used at 1:1000 and manufacturer information can be found in the Key Resources table. HRP-conjugated secondaries are used at 1:5000 (BioRad).

### RNA-sequencing and bioinformatics analysis

cDNA libraries were constructed from tissue-extracted RNA samples according to the manufacturer’s protocol (Roche KAPA HyperPrep kit). Libraries were sequenced on the NovaSeq6000 platform (Novogene). Transcriptome mapping of RNA-seq reads (20-30 million reads per sample) was performed with STAR^74^ using the Ensembl annotation of the mm10 reference genome. Read counts for individual genes were generated using HTSeq.^75^ Differential expression analysis was performed using EdgeR^76^ after normalizing read counts and including only those genes with CPM > 1 for at least one sample. Differentially expressed genes (DEGs) were defined based on the criteria of >2-fold absolute change in expression value and adjusted P-value (FDR) < 0.05 for Heart and DMD; and >1.5-fold change and FDR<0.05 for Sarcopenia. Analysis of enriched functional gene categories was performed using Gene Set Enrichment Analysis (GSEA).^77^

### MicroRNA Expression Analysis

For mouse tissues, miRNA and total RNA was purified from previously flash frozen tissue using the miRNeasy kit (Qiagen) according to the manufacturer’s instructions. Reverse transcription (RT) was then performed using the TaqMan MicroRNA Reverse Transcription Kit (Applied Biosystems) with RT probes for U6 and miR-128-3p. For miR-27a, reverse transcription was performed using the TaqMan Advanced miRNA Assay (Applied Biosystems). miR-128-3p (or miR-27a) and U6 expression was quantified by real-time PCR using TaqMan Fast Advanced Master Mix (Applied Biosystems) using the QuantStudio6 Real-Time PCR system (Applied Biosystems). miR-128-3p (or miR-27a) Ct values were normalized to U6 Ct values measured in the same run. RT and TaqMan probes were purchased from Applied Biosystems as part of the TaqMan microRNA Assay.

For dog serum, microRNAs were isolated from frozen serum samples (100-200 µL per sample point) using the miRNeasy serum/plasma kit (Qiagen, #217184). cDNA was prepared via miScript II RT kit (Qiagen, #218161) using 3 µL of serum RNA. All cDNA preparations were subsequently diluted 1/20 with nuclease free water. Following cDNA synthesis, qPCR was conducted via the miScript PCR System (Qiagen, #218073) in 10 µL volumes (2 µL cDNA) using a CFX384 light-cycler (BioRad), with primers specific to *Canis familiaris* miR-128-3p (Qiagen, #MS00029449) or to miR-223 (#MS00030126, a reference microRNA for canine serum,^78^ along with a universal primer (Qiagen, #218073). Melt curves were included as standard. After conversion of Cq to relative quantity (RQ = efficiency ^(Cq^ ^min–Cq^ ^sample)^), miR-128-3p data was normalized to miR-223 to obtain relative expression. Expression values were normalized via log transformation for statistical analysis.

For dog muscle, frozen muscle samples were crushed with pestle and mortar that were pre-chilled on dry ice and collected into a 1.5 mL Eppendorf. MicroRNAs were extracted using a miRNeasy kit (Qiagen, #217084), according to manufacturer’s instructions. RNA concentration and purity were analyzed using a NanoDrop 1000 spectrophotometer (Thermo Fisher Scientific). Samples with a 260 nm:230 nm absorbance ratio of less than 1.7 underwent a further isopropanol precipitation step. MicroRNAs were reverse transcribed using a miRCURY LNA RT kit (Qiagen, #339340) with the addition of the provided UniSp6 RNA spike-in control and the cel-miR-39-3p RNA spike-in template (Qiagen, #339390), according to manufacturer’s instructions. Samples were diluted to a concentration of 5 ng/µL in nuclease free H_2_O and reverse transcribed using 10 ng total RNA per 10 µL reaction. cDNA was used in qPCR assays with primers (Qiagen, #339306) to our validated panel of 4 reference RNAs (NF4)^79^: cfa-miR-15a (YP02103582), hsa-let-7b-5p (YP00204750), cfa-miR-125a (YP00203952) and cfa-miR-191 (YP00205972) and to hsa-miR-128-3p (YP00205995) with additional spike in controls cel-miR-39-3p (YP00203952) and UniSp6 (YP00203954). qPCR was performed using the miRCURY LNA SYBR Green PCR Kit (Qiagen, #339347), according to the manufacturer. Inter-plate calibration was achieved by using a single plate to quantify a subset of microRNAs in samples from each original plate (N=4 common miRCURY assays per plate); the mean difference in Cq value was determined for each plate and was subtracted from the original Cq result to give the adjusted mean Cq.

For pig tissues, three snap-frozen myocardium tissue cubes with an edge length of approximately 4 mm were homogenized together for each animal. Tissue samples stored at −80°C were wrapped in aluminium foil and cooled to −196°C in liquid nitrogen. The samples were then pulverized while taking care to avoid warming of the tissue. The resulting powder was further ground in a mortar placed in liquid nitrogen. Finally, 25 mg of the powder was added to 700 μl QIAzol lysis reagent and directly homogenized using a Polytron PT 2500 E rotor-stator homogenizer. Total RNA was extracted using the miRNeasy kit (Qiagen) according to the manufacturer’s protocol. Reverse transcription was performed using the TaqMan MicroRNA Reverse Transcription Kit (Applied Biosystems), together with miR-128-3p and U6 control assays (Applied Biosystems). Quantitative PCR was carried out using TaqMan Fast Advanced Master Mix (Applied Biosystems) and the corresponding miR-128-3p and U6 control assays (Applied Biosystems). Real-time polymerase chain reaction was performed on a Roche LightCycler® 96 instrument.

For patient serum, miRNA was extracted using the *mir*Vana PARIS RNA and Native Protein Purification Kit (Invitrogen) according to the manufacturer’s instructions. Reverse transcription was then performed using the TaqMan Advanced miRNA Assay (Applied Biosystems). miR-128-3p and the reference miRNA, miR-93, expression was quantified by real-time PCR using TaqMan Fast Advanced Master Mix (Applied Biosystems) using the QuantStudio6 Real-Time PCR system (Applied Biosystems). miR-128-3p Ct values were normalized to miR-93 Ct values measured in the same sample, and relative expression was calculated using the ΔΔCt method. TaqMan probes were purchased from Applied Biosystems as part of the TaqMan microRNA Assay.

### Sample Preparation for SplintR Ligation qPCR

We performed sample preparation as previously described by Shin et al. with modifications.^80^ In brief, tissues, including liver, heart, kidney, gastrocnemius (GA), and tibialis anterior (TA), were harvested and homogenized in 300–1000 μL of tissue lysis buffer (10 mM Tris-HCl [pH 7.5], 150 mM NaCl, 1% NP-40, 0.5% sodium deoxycholate, 0.1% sodium dodecyl sulfate, 5 mM ethylenediaminetetraacetic acid, and 1 mM ethylene glycol-bis(β-aminoethyl ether)-N,N,N′,N′-tetraacetic acid). Homogenates were incubated on ice for 1 hour with gentle vortexing, followed by centrifugation at 17,500 × g for 30 minutes at 4 °C. The supernatants were collected, and protein concentrations were determined using the Pierce BCA Protein Assay (Thermo Fisher Scientific) according to the manufacturer’s instructions. Tissue lysates from the same tissue type were normalized to equal protein concentrations (2–20 mg/mL) using tissue lysis buffer. Blood and the indicated tissues were also collected from an untreated mouse and processed in parallel using the same protocol to serve as blank controls and as matrix diluents for the standard curve in SplintR ligation qPCR. Blood from the untreated mouse was collected via cardiac puncture to ensure sufficient serum volume for downstream applications.

### Quantification of anti-miR-128 ASO by SplintR Ligation-qPCR

We performed SplintR Ligation-qPCR as previously described by Shin et al. with modifications.^80^ In brief, standard anti-miR-128 ASO samples (1 μM, 0.1 μM, 10 nM, 1 nM, 0.1 nM, and 10 pM) were prepared in serum or the indicated tissue lysates from untreated mice. The SplintR ligation reaction was assembled in a total volume of 12 μL, consisting of 2 μL of 10× SplintR ligase reaction buffer (New England Biolabs), 2 μL of probe A (100 nM), 2 μL of probe B (100 nM), 2 μL of either standard (1 μM to 10 pM) or sample, and nuclease-free water. Probe–target hybridization was performed by heating the reaction mixture to 95°C for 10 minutes, followed by gradual cooling to 37°C at a rate of 0.1°C per second. After hybridization, 8 μL of diluted SplintR ligase (2.5 U per reaction) was added to each mixture. Ligation was carried out at 37°C for 30 minutes, followed by enzyme inactivation at 65°C for 20 minutes. For qPCR quantification of ligated products, reactions were prepared in a final volume of 10 μL containing 2 μL of the ligation mixture, 5 μL of TaqMan™ Fast Advanced Master Mix (Thermo Fisher Scientific), 0.5 μL of a primer/probe mix (forward primer, reverse primer, and TaqMan probe at 10 μM, 10 μM, and 5 μM, respectively), and nuclease-free water. Quantitative PCR was performed in duplicate using the QuantStudio™ 6 system (Applied Biosystems) under the following cycling conditions: initial denaturation at 95°C for 20 seconds, followed by 40 cycles of 95°C for 5 seconds and 60°C for 10 seconds. The probes and primers were synthesized by Integrated DNA Technologies are listed below:

Probe A: 5’-AGCTCGACCTCTCTATGGGCAGTCACGACAGCACAGTG-3’

Probe B: 5’-/5Phos/AACCGGTCGCTGAGTCGGAGACACGCAGGGCTTAATG-3’

Forward: 5’-AGCTCGACCTCTCTATGGG-3’

Reverse: 5’-CATTAAGCCCTGCGTGTCT-3’

TaqMan Probe: 5’-/56-FAM/AGTCACGAC/ZEN/AGCACAGTGAACCG/3IABkFQ/-3’

### H&E Staining

For mouse toxicity studies, liver and kidney tissues were dissected, fixed in 4% paraformaldehyde for 48 hours, and transferred to 70% ethanol. Paraffin embedding, sectioning, and H&E staining was performed at the UC San Francisco Liver Center. For monkey toxicity studies, tissue preparation and staining was performed at the Pathology Lab at Hvidovre Hospital, Copenhagen.

### Mouse Skeletal Muscle Fibrosis Quantification

Tibialis anterior muscle was embedded in OCT, cryosectioned from the middle of the tissue into cross sections at 10 µm, and adhered to slides. Sections were fixed on slides in formalin for 1 hour then dehydrated in 70%, 95%, 100% ethanol. Slides were incubated in Bouin’s fixative for 1 hour at 55°C or overnight at room temperature. Next, slides were washed in tap water until yellow was cleared, incubated 20-25 min in 0.1% fast green in distilled water, 2 min in 1% acetic acid, washed with tap water, 30 min in 0.1% Sirius Red (Sigma) in saturated picric acid, washed again with tap water, and dehydrated 30 sec each in 95% and 100% ethanol. Finally, they were incubated 30 seconds in xylenes before cover slipping. Tissue was imaged using a Zeiss AxioImager microscope and staining was quantified using ImageJ.

### Mouse Biochemical Assays (ALT, AST, BUN, Creatinine, Creatine Kinase)

Serum metabolites were measured by assay kits according to the manufacturer’s protocol (see Key Resources table).

### Non-human Primate Chemistry Panels

Serum chemistry panels including alanine aminotransferase (ALT), aspartate aminotransferase (AST), blood urea nitrogen (BUN), and creatinine, as well as complete blood counts, were performed by Antech Diagnostics at each blood collection timepoint to assess hepatic and renal function.

### Dog Blood collection

Blood was obtained at monthly intervals (1-18 months of age) by jugular venipuncture into plain tubes. After clotting, blood samples were centrifuged (Heraeus, #3328) at 500 x g for 10 min at 4°C. Serum was collected and frozen at -80°C until processing. 208 samples in total were used (WT: N=96; DE50: N=112), with N≥3 per genotype per timepoint (mean: 5-6) for all except month 1.

### Dog muscle tissue collection

Longitudinal vastus lateralis biopsy samples were obtained from 6 male WT and 6 male DE50-MD dogs, as described in detail elsewhere.^79^ Briefly, dogs were pre-medicated with intravenous methadone (0.2 mg/kg, Comfortan, Dechra) via venous catheter, prior to induction of anesthesia by intravenous propofol (1-4 mg/kg, PropoFlo Plus, Zoetis). Anesthesia was maintained by sevoflurane (1.5-3.5%, SevoFlo, Zoetis) in oxygen via endotracheal tube using a Aestiva/5 anesthesia machine (Datex-Ohmeda). Vastus lateralis muscle samples were biopsied at 3-monthly intervals between 3 and 18 months of age. A panel of muscles was also collected post-mortem in 3 dogs of each genotype that were euthanized at the end of the 18-month study period. Four muscles were collected post-mortem: cranial tibial (CRT), semimembranosus (SEM), lateral head of the triceps (TRI) and the diaphragm (DIA). The longitudinal and post-mortem sets were non-overlapping for WT; one DE50-MD dog contributed to both sets, yielding N = 9 unique WT and N = 8 unique DE50-MD dogs overall. Muscle samples were unavailable for one DE50-MD dog at the 15-month VL collection and one WT dog at the 18-month VL collection; therefore, the complete dataset comprised 70 longitudinal VL biopsy samples (35 per genotype) and 24 post-mortem muscle samples (12 per genotype), yielding 47 muscle samples per genotype. Samples were collected into cryovials and snap-frozen in liquid nitrogen, before transfer to -80°C storage until use.

During the biopsy procedure, dogs were administered intravenous cefuroxime antibiotic (20 mg/kg, Zinacef, GSK) and carprofen for analgesia (2 mg/kg, Rimadyl, Pfizer). Following the procedure, post-operative carprofen (2 mg/kg, Rimadyl, Pfizer) was given orally, once daily for 3 days.

### Pig Blood collection and clinical chemistry

Blood collection took place from the right jugular vein, using Serum Monovettes (Sarstedt, Nümbrecht, Germany). Serum Monovettes were kept at room temperature for 30 min for clotting, followed by centrifugation at 1800 g for 20 min at 4°C. Serum was aliquoted and stored at −80°C until further processing (storage period did not exceed 6 months). Troponin I levels were measured by the CMIA method (SYNLAB vet GmbH, Augsburg, Germany).

### Echocardiography

*(mouse and pigs)* For mice, serial transthoracic echocardiography was performed in mice anesthetized with 1.5% isoflurane at baseline (pre-MI) and 2- or 3-days post-MI, as well as at 7, 14, 21, and 28 days post-MI using a Vevo 3100LT high-resolution micro-ultrasound system (VisualSonics Inc., Toronto, Canada). Two-dimensional (B-mode) images were acquired in parasternal long-axis and short-axis views. Left ventricular (LV) end-systolic volume (ESV) and end-diastolic volume (EDV) were measured using Simpson’s method with VevoLab software (VisualSonics Inc.), and ejection fraction (EF) was calculated accordingly. LV anterior and posterior wall thickness at end-diastole were measured using M-mode imaging at apical, mid-ventricular, and basal levels.^70^ Echocardiograph acquisition and all analyzes were performed by an investigator blinded to treatment allocation, including anti-miR-128 administration.

For pig echocardiography, pigs were sedated with 20 mg/kg ketamine (Ursotamin, Serumwerke Bernburg, Bernburg, Germany) and 2 mg/kg azaperone (Azaporc, Serumwerke Bernburg), followed by an anesthesia with 4 mg/kg/h of propofol (Propofol 2%, Fresenius Kabi, Bad Homburg, Germany). After sufficient depth of anesthesia has been achieved, pigs were placed in right lateral recumbency, and heart function was measured by standard 2D transthoracic echocardiography (Esaote MyLab X8) as previously described.^45,81^ Left ventricular ejection fraction and fractional shortening were determined by M-mode method. All measurements were performed by the same investigator, who was blinded to the treatment and placebo groups.

### Quantification of Infarct Scar Size

Mice were euthanized 28 days after MI, and hearts were arrested in diastole by saturated KCl injection, embedded in O.C.T. compound (Sakura Finetek, USA), and frozen for cryosectioning. Heart was sectioned transversely at 10-μm thickness with 300-μm intervals between sections. Eight sections spanning from the ventricular apex to the base were collected and stained with Picrosirius Red to quantify fibrotic collagen (red) deposition as follows: Heart sections were first incubated for 1 hour in Solution I (0.01% Fast Green FCF in saturated picric acid; Sigma-Aldrich, F7252 and P6744), followed by 1 hour in Solution II (0.04% Fast Green FCF and 0.1% Sirius Red in saturated picric acid; Sigma-Aldrich, 365548). Sections were rinsed in acidified water, dehydrated in 100% ethanol, cleared in xylene (Avantor), and mounted for imaging and analysis.

Quantitative analysis of fibrotic collagen was performed on images acquired using a 1x objective and analyzed with ImageJ. Infarct scar size was measured by investigators blinded to treatment groups using two established approaches, depending on the MI model. For the I/R MI model, scar area and total LV area were measured in each section. Infarct scar size was expressed as a percentage of the LV by dividing the sum of infarct areas from all sections by the sum of LV areas from all sections and multiplying by 100, as previously described.^69^ For the permanent LAD ligation MI model, infarct scar size was quantified using the midline arc length method, as previously described.^82^

### Quantification of Non-Scar Fibrosis

In the permanent LAD ligation MI model, non-scar fibrosis was quantified from images acquired with a 40x objective. For each heart, the cryosection containing the largest infarct area, identified from Picrosirius Red-stained sections, was selected for analysis. Fibrosis near the infarct scar zone and the peri-infarct border zone was measured using ImageJ by an investigator blinded to treatment groups, as previously described.^69^

### Immunofluorescence staining

Tibialis anterior or gastrocnemius muscle was embedded in OCT, cryosectioned from the middle of the tissue into cross sections at 10 µm and adhered to slides. To begin staining, slides were warmed to room temperature for 30 minutes. Then, they were washed three times for 5 mins in PBS, permeabilized in 0.4% Triton X-100 in PBS for 10 minutes, and blocked for 2 hours in 5% normal goat serum (Invitrogen) in 0.05% Triton X-100 in PBS (PBS-T) at room temperature. Sections were incubated overnight with primary antibodies in 5% normal goat serum in 0.05% PBS-T at 4°C. The next day, sections were washed three times for 5 minutes in 0.05% PBS-T. Secondary antibody incubation was then performed for 2 hours at room temperature in 5% normal goat serum in 0.05% PBS-T. Finally, slides were washed three times for 5 minutes in 0.05% PBS-T. Slides were coverslipped with Prolong Antifade Gold Mountant (Invitrogen). Slides were observed and imaged via Zeiss Axio Imager M2 or Zeiss Spinning Disk Confocal. Primary antibody manufacturers and catalog numbers can be found in the Key Resources table. Primary antibodies were used at the following concentrations: Laminin (1:200) and MyHC I/IIa/IIb (3 µg/mL).

### Muscle Fiber Cross Sectional Area Quantification

Sectioned muscles underwent immunofluorescent staining using an Anti-Laminin primary antibody (Sigma) and Alexa-Fluor 488 secondary antibody. Entire tissue sections were imaged using a Zeiss spinning disc confocal microscope, and muscle fiber cross sectional area was quantified in ImageJ.

### Transmission Electron Microscopy

Electron microscopy was performed at the UC Berkeley Electron Microscope Laboratory. Samples were fixed in 2% paraformaldehyde 2.5% glutaraldehyde in 0.1M sodium cacodylate buffer, pH 7.4. For processing, samples were rinsed 3x 5min in 0.1M sodium cacodylate buffer, pH 7.2, put in 1% osmium tetroxide in 1.6% potassium ferricyanide for 1-2 hours, rinsed 3x 5min in 0.1M sodium cacodylate buffer, pH 7.2, rinsed 5 min in distilled water, and dehydrated in acetone (35%, 50%, 70%, 80%, 95%, 100%, 100%; 10 min each). Samples were then infiltrated with 2:1 acetone: resin (with accelerator-BDMA) for 1 hour, 1:1 acetone: resin for 1 hour, and 1:2 acetone: resin overnight. The next day, samples were put in 100% resin (with accelerator) 3x 1-2 hours. Samples were embedded in molds and baked at > 60°C for 2 days. Samples were then visualized via the TECNAI 12 TEM.

### Mitochondrial DNA Copy Number

Muscle tissue was dissected and snap frozen in liquid nitrogen. DNA was extracted using the DNeasy Blood and Tissue Kit (Qiagen) according to the manufacturer’s instructions. Mitochondrial and nuclear DNA copy number were quantified by measuring CoxIII and beta-actin respectively using PowerUP SYBR Green (Applied Biosystems) using the QuantStudio6 Real-Time PCR system (Applied Biosystems). CoxIII Ct values were normalized to beta-actin levels using the ΔΔCt calculation.

### Proteomics and Bioinformatics Analysis

Heart and skeletal muscle tissue samples were snap-frozen in liquid nitrogen and cryo-pulverized using a CP02 Automated Dry Pulverizer (Covaris, Woburn, MA, USA) according to the manufacturer’s instructions. Tissue powder was lysed in 8 M urea/0.5 M NH4HCO3 by ultrasonication (18 cycles of 10 s) using a Sonopuls HD3200 (Bandelin, Berlin, Germany). Proteins were quantified with Pierce 660 nm Protein Assay (Thermo Fisher Scientific, Rockford, IL, USA). Twenty micrograms of protein were digested with Lys-C (FUJIFILM Wako Chemicals Europe GmbH, Neuss, Germany) for 4 h and subsequently with modified porcine trypsin (Promega, Madison, WI, USA) for 16 h at 37°C.

One microgram of the digest was injected on an UltiMate 3000 nano-LC system coupled online to a Q-Exactive HF-X instrument (Thermo Fisher Scientific). Samples were transferred to a PepMap 100 C18 trap column (100 µm×2 cm, 5 µM particles, Thermo Fisher Scientific) and separated on an analytical column (PepMap RSLC C18, 75 µm×50 cm, 2 µm particles, Thermo Fisher Scientific) at 250 nL/min with an 80-min gradient of 5-20% of solvent B followed by a 9-min raise to 40%. Solvent A consisted of 0.1% formic acid in water and solvent B of 0.1% formic acid in acetonitrile. MS spectra were acquired using a top 15 data-dependent acquisition method on a Q Exactive HF-X mass spectrometer. Raw files were processed with Maxquant (v.1.6.7.0),^83^ using its built-in search engine Andromeda and the NCBI RefSeq Sus scrofa database (v.7-5-2020), applying the default search parameters recommended for Orbitrap mass spectrometers. Protein intensities were normalized using the MaxLFQ approach.^84^ Statistical analysis and visualization were done using Perseus and R framework. Proteins detected in all replicates of at least one condition were kept for quantitative analysis. Missing values were imputed by Perseus default parameters. The volcano plots were generated using two-tailed Student’s t-test and permutation-based FDR cut-off of 0.05, together with an s0-parameter of 0.1 to additionally consider fold changes. The mass spectrometry proteomics data have been deposited to the ProteomeXchange Consortium via the PRIDE^85^ partner repository with the dataset identifier PXD082523.

### Statistical Analysis

Statistical details are reported in each figure legend. Data are represented as mean values, and error bars represent SD. Data analysis was performed using GraphPad Prism 10 or Stata 13.1 (for two-way ANOVA calculations). For mouse echocardiographic parameters measured across multiple groups and time points, a two-factor repeated-measures analysis of variance (ANOVA) was performed using a mixed-effects model with restricted maximum likelihood estimation and an unstructured residual covariance matrix. Differences over time and between treatment conditions were evaluated using contrasts and pairwise comparisons with Šidák adjustment for multiple testing. Porcine ejection fraction (EF) and fractional shortening (FS) data were analyzed using linear mixed-effects models with group (anti-miR-128, scramble), time (baseline, 2 months), and the group-by-time interaction as fixed effects and animal as a random effect (PROC MIXED, SAS). Degrees of freedom were estimated using the Kenward-Roger method. Canine serum and muscle miR-128-3p expression were compared between DE50-MD and wildtype dogs using linear mixed models on log-transformed relative quantities, with genotype, sampling occasion (age in months for serum; muscle and age group for muscle) and the genotype by sampling-occasion interaction as fixed effects, and dog as a random effect, accounting for repeated sampling of individual animals across timepoints and for multiple muscles sampled from the same animal. Denominator degrees of freedom were estimated by the Satterthwaite approximation, and genotype comparisons at individual sampling occasions were derived from estimated marginal means without adjustment for multiple comparisons (SPSS Statistics). All mouse experiments were repeated at least twice. Large animal studies were only conducted once in accordance with ethical guidelines and best practices. Single studies were sufficient as results were concordant with previous mouse studies.

