## Supplementary Figures and Tables for "A positively selected microRNA controls a reversible aging program in striated muscle"

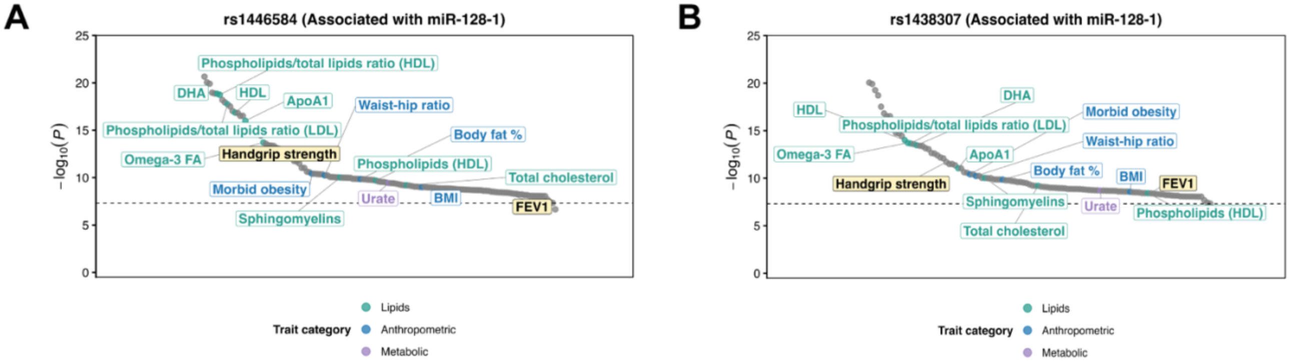


**Figure S1. Additional SNPs in the miR-128-1 locus associate with muscle and pulmonary function in humans.**

(A) PheWAS plot of the single nucleotide polymorphism (SNP) rs1446584 at the miR-128-1 locus. Dashed line denotes genome-wide significance ($P<5 \times 10^{-8}$).
(B) PheWAS plot of the single nucleotide polymorphism (SNP) rs1438307 at the miR-128-1 locus. Dashed line denotes genome-wide significance ($P<5 \times 10^{-8}$).

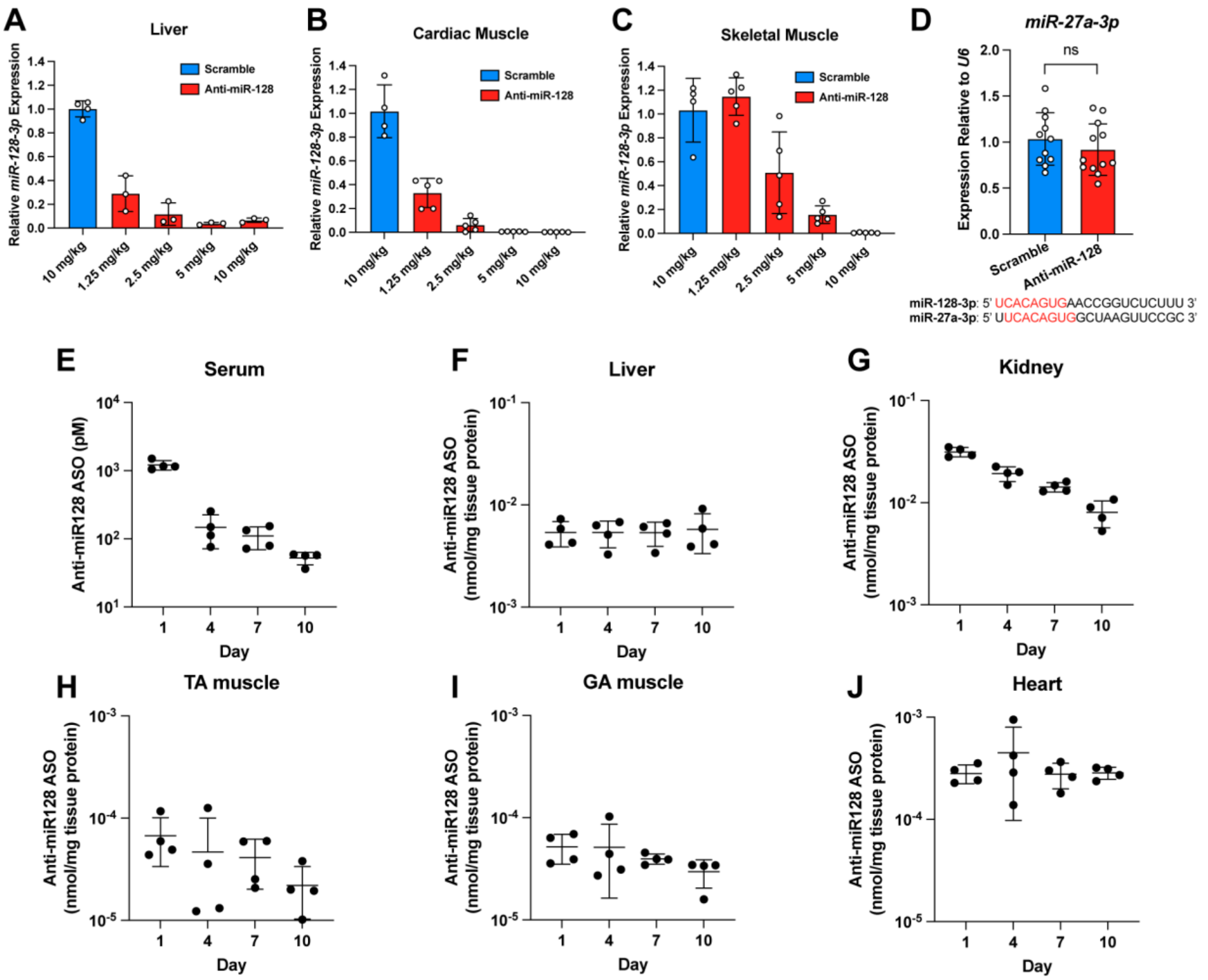


**Figure S2. Dose response, specificity, and tissue kinetics of anti-miR-128.**

(A-C) *miR-128-3p* expression levels measured 28 days after a single treatment of C57BL/6J mice with scramble (10 mg/kg) or anti-miR-128 (10, 5, 2.5, 1.25 mg/kg) in liver (A), cardiac muscle (B), or skeletal muscle (C).
(D) RT-qPCR analysis of non-target microRNA, miR-27a-3p following anti-miR-128 or scramble treatment in gastrocnemius muscle of 106-week-old C57BL/6J mice that had been treated for 20 weeks. The sequence comparison of miR-128-3p to miR-27a-3p is shown, with shared nucleotides highlighted in red.
(E-J) Anti-miR-128 concentrations measured by SplintR qPCR at 1, 4, 7, or 10 days after a single treatment at 10 mg/kg in C57BL/6J mice in serum (E), liver (F), kidney (G), TA muscle (H), GA muscle (I), and heart tissue (J).

Data are presented as mean +/- SD. N=4 per group (A-C, E-J) and N=11-12 per group (D). ns= not significant (unpaired two-tailed t-test (D)).

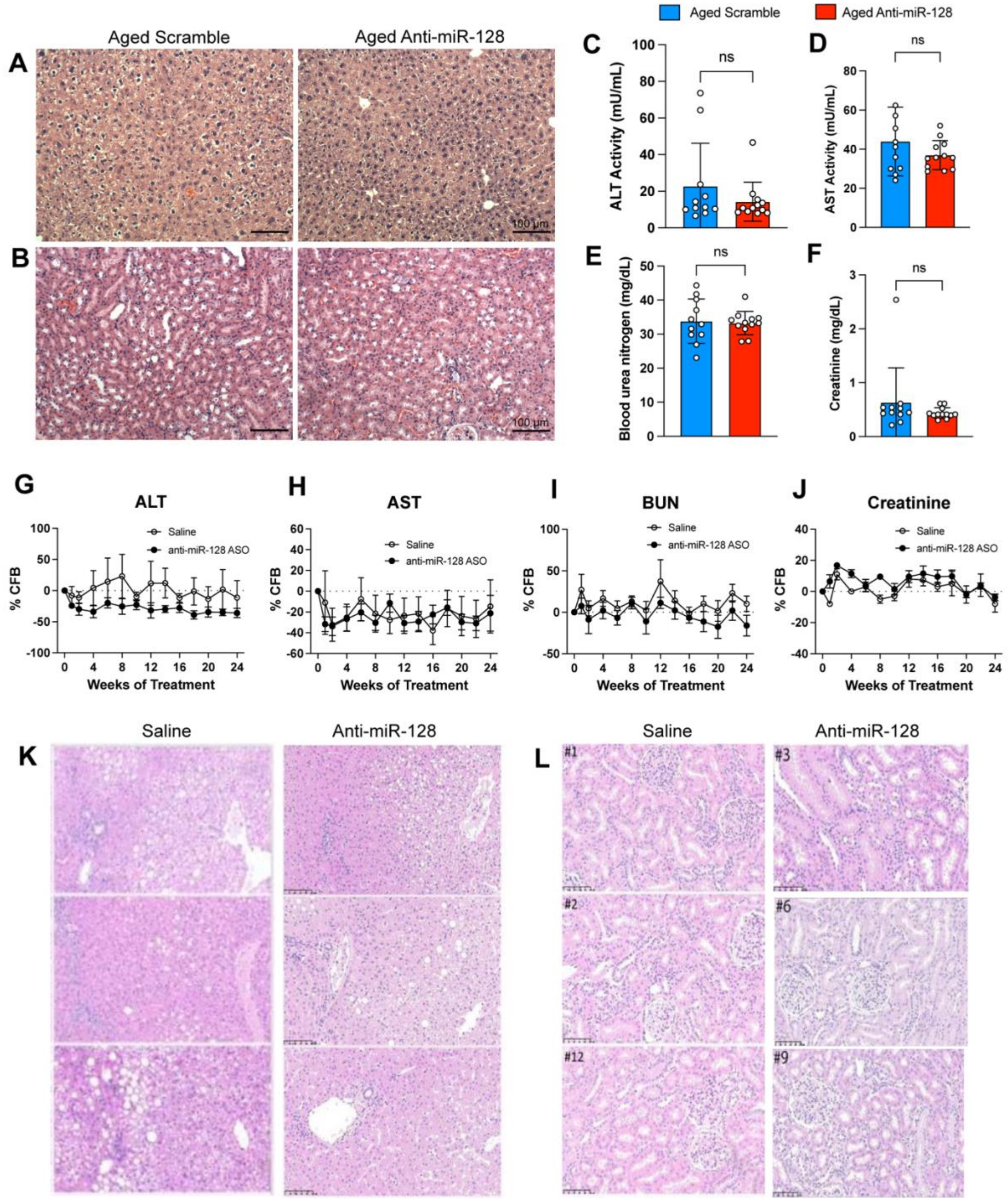


**Figure S3. Cross-species safety profile of long-term anti-miR-128 treatment.**

(A-F) Toxicity analysis of serum and tissue from 106-week-old C57BL/6J mice that had been treated with anti-miR-128 or scramble at 10 mg/kg for 20 weeks. H&E-stained liver (A) and kidney (B), scale bar, 100 µm (inset). Serum ALT activity (C), AST activity (D), blood urea nitrogen (E), and creatinine (F) measured at 20 weeks of treatment.

(G-L) Toxicity analysis of serum and tissue from cynomolgus monkeys treated with saline or anti-miR-128 at 5 mg/kg for 24 weeks. Serum ALT activity (G), AST activity (H), blood urea nitrogen (I), and creatinine (J) measured at 14 scheduled timepoints during treatment. H&E-stained liver (K) and kidney (L) tissue, scale is 100 μm (inset).

Data are presented as mean +/- SD. N=11-12 per group (mice); N=3-4 per group (monkeys) ns, not significant by unpaired, two-tailed, t-test (C-F).

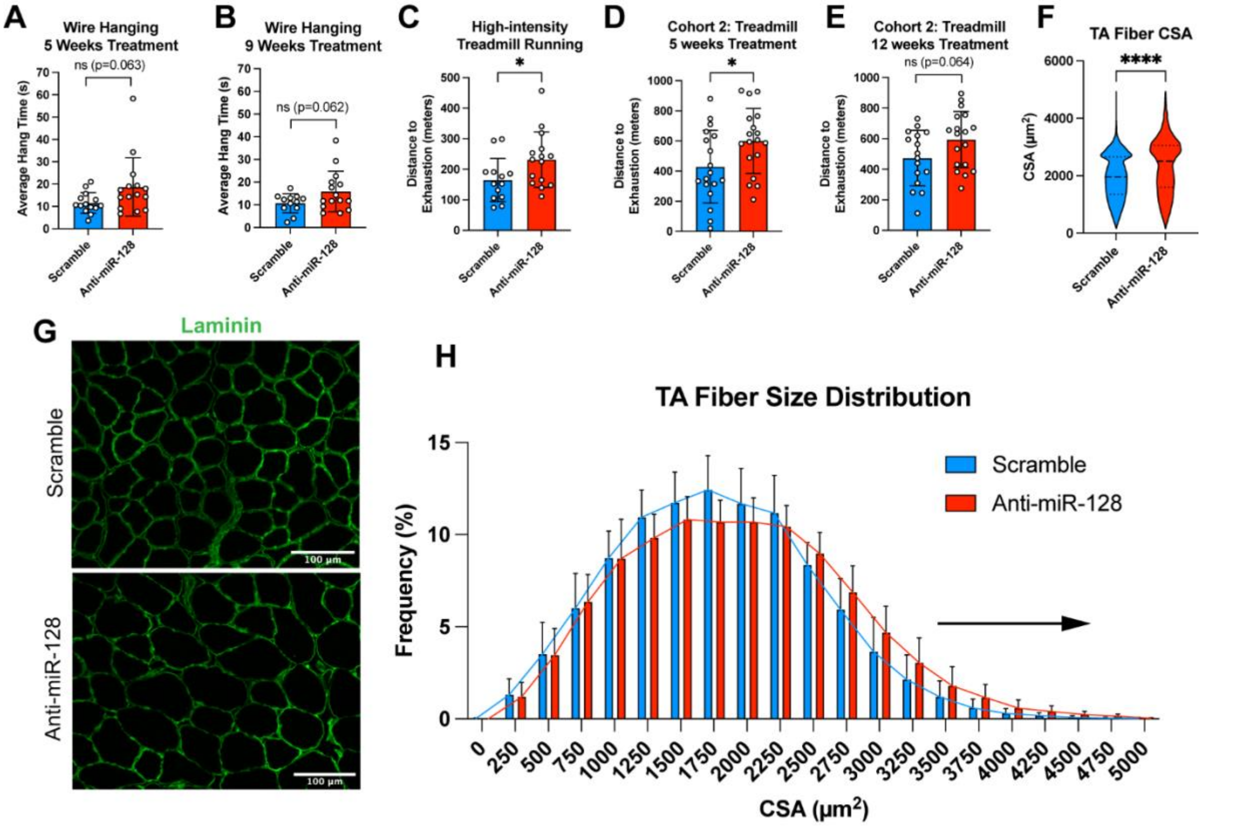


**Figure S4. Additional functional and histological analyses in aged mice treated with anti-miR-128.**

(A) Wire hanging at 5 weeks of treatment from cohort 1.
(B) Wire hanging at 9 weeks of treatment from cohort 1.
(C) Distance run to exhaustion using a high intensity treadmill protocol at 8 weeks of treatment from cohort 1.
(D) Distance run to exhaustion at 5 weeks of treatment from cohort 2.
(E) Distance run to exhaustion at 12 weeks of treatment from cohort 2.
(F-H) Muscle fiber cross sectional area (CSA) by laminin staining from TA muscle of scramble or anti-miR-128 treated mice from cohort 1. Fiber CSA by group (F). Representative immunofluorescence images of TA muscle stained for laminin (Scale bar, 100 µm; inset) (G). Frequency distribution of TA muscle fiber CSA quantified for each mouse and averaged by treatment group (H).

Data are presented as mean +/- SD. N=14-15 per group (A); N=12-15 per group (B); N= 13-15 per group (C); N=18 (D); N=16-18 (E); N=11 per group (H). ns, not significant, *p<0.05,**p<0.01,***p<0.001,****p<0.0001 (unpaired, two-tailed t-test (A-E), Kolmogorov-Smirnov test (F)).

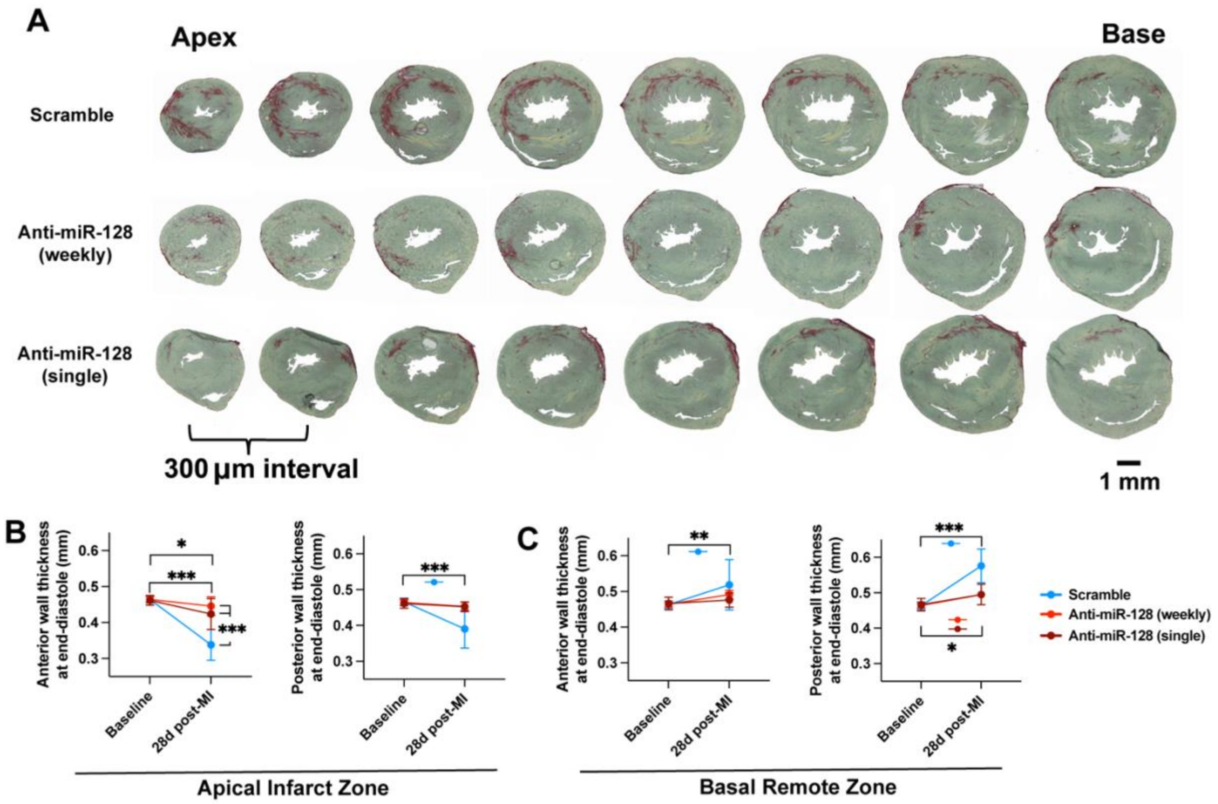


**Figure S5. Additional fibrosis images and wall thickness measurements from post-I/R mice.**

(A) Representative serial cryosections from the apex to base of the ventricle for infarct scar size measurement after I/R MI (cohort 1). Heart samples were sliced transversely at 10-µm thickness with an interval of 300 µm between each section. Eight cryosections from the apex to base of the ventricle were stained with Picrosirius Red and Fast Green for infarct scar size measurement. Mice were treated with scramble control ASO (weekly injections; top), or anti-miR-128 (weekly injections; middle), or anti-miR-128 (single injection; bottom).
(B-C) Anterior and posterior wall thickness by echocardiography at apical infarct zone (B) and basal remote zone (C) at baseline and 28 days post-I/R from cohort 1.

Data are presented as mean +/- SD. N=8-10 per group (B-C). *p<0.05, **p<0.01,***p<0.001,****p<0.0001 (two-way ANOVA followed by the Šidák method (B-C)).

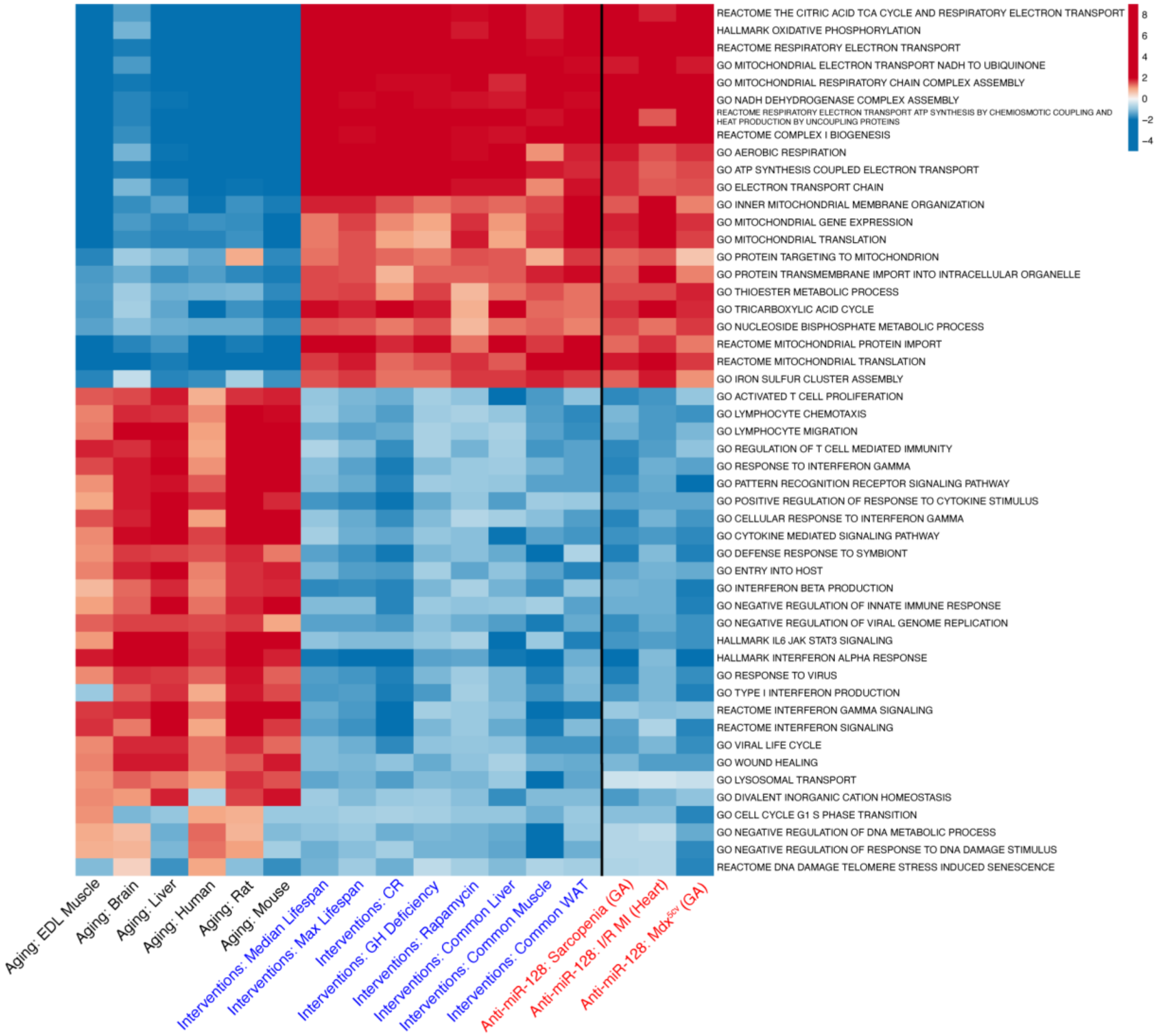


**Figure S6. Expanded pathway analysis of mitochondrial and inflammatory transcriptional signatures rescued by anti-miR-128 across aging and disease contexts.**

Gene set enrichment analysis of the expression profile of aged C57BL/6J gastrocnemius muscle, *mdx*$^{5cv}$ gastrocnemius muscle, and post-I/R cardiac tissue treated with anti-miR-128 (red text) compared to signatures of mammalian aging (black text) and signatures of longevity interventions (blue text). Red boxes indicate upregulated pathways, and blue boxes indicate downregulated pathways. Additional pathways within the mitochondrial and inflammatory categories are shown beyond those depicted in Figure 7.

### Supplementary Table 1. Serum Chemistry & Hematology — NHP Safety Study

Data are presented as mean ± SD. N = 3 (saline), 4 (anti-miR-128) per group. ns, not significant; *p<0.05, **p<0.01, ***p<0.001 (unpaired, two-tailed Welch's t-test vs. saline at the same timepoint). The primary analysis reported here is change from each animal's own week-0 value (see Results); week-0 comparisons precede the first dose and reflect baseline group differences, not treatment effects. Reference intervals have been omitted: those supplied with the source panels are generic adult-primate ranges rather than cynomolgus-specific, and several analytes fall wholly outside them in both arms at every timepoint, amylase most clearly (observed 92-248 U/L against a reported interval of 1,000-2,500 U/L). CPK, see footnote a; chloride, see footnote b.

| Analyte | Units | Saline | | | | Anti-miR-128 | | | |
|---|---|---|---|---|---|---|---|---|---|
| | | Week 0 | Week 8 | Week 16 | Week 24 | Week 0 | Week 8 | Week 16 | Week 24 |
| **HEPATIC FUNCTION** | | | | | | | | | |
| Total protein | g/dL | 7.3 ± 0.5 | 7.3 ± 0.4 | 7.4 ± 0.3 | 7.8 ± 0.6 | 7.9 ± 0.2 | 7.7 ± 0.2 | 7.7 ± 0.2 | 8.0 ± 0.5 |
| Albumin | g/dL | 3.6 ± 0.2 | 3.7 ± 0.1 | 3.8 ± 0.1 | 3.8 ± 0.2 | 3.8 ± 0.1 | 3.8 ± 0.2 | 3.8 ± 0.1 | 3.7 ± 0.1 |
| Globulin | g/dL | 3.7 ± 0.3 | 3.6 ± 0.3 | 3.6 ± 0.2 | 4.0 ± 0.6 | 4.1 ± 0.2 | 3.9 ± 0.2 | 3.9 ± 0.2 | 4.3 ± 0.4 |
| AST | U/L | 61.0 ± 23.9 | 43.7 ± 15.0 | 34.0 ± 5.3 | 45.3 ± 15.0 | 43.2 ± 6.2 | 29.5 ± 4.8 | 34.2 ± 12.7 | 33.0 ± 13.3 |
| ALT | U/L | 64.0 ± 21.3 | 77.0 ± 37.6 | 56.0 ± 16.1 | 50.3 ± 18.8 | 31.2 ± 8.3 | 22.5 ± 5.9 | 22.8 ± 8.5 | 20.0 ± 6.8 |
| Alkaline phosphatase | U/L | 209.0 ± 67.5 | 182.7 ± 83.6 | 146.0 ± 58.6 | 158.0 ± 59.4 | 144.8 ± 33.5 | 138.0 ± 34.2 | 110.8 ± 26.4 | 125.8 ± 33.5 |
| GGT | U/L | 98.3 ± 24.6 | 94.3 ± 25.0 | 99.3 ± 28.2 | 91.3 ± 26.3 | 99.5 ± 34.6 | 96.5 ± 34.2 | 82.8 ± 27.0 | 81.5 ± 26.1 |
| Total bilirubin | mg/dL | 0.2 ± 0.1 | 0.1 ± 0.1 | 0.2 ± 0.1 | 0.2 ± 0.1 | 0.2 ± 0.1 | 0.1 ± 0.0 | 0.1 ± 0.0 | 0.1 ± 0.1 |
| **RENAL FUNCTION** | | | | | | | | | |
| BUN | mg/dL | 11.3 ± 2.1 | 12.7 ± 1.5 | 11.0 ± 1.0 | 12.3 ± 1.5 | 12.5 ± 1.9 | 13.8 ± 1.3 | 11.5 ± 0.6 | 10.2 ± 2.2 |
| Creatinine | mg/dL | 1.3 ± 0.3 | 1.2 ± 0.3 | 1.3 ± 0.3 | 1.2 ± 0.3 | 1.3 ± 0.2 | 1.5 ± 0.2 | 1.5 ± 0.2 | 1.3 ± 0.1 |
| Phosphorus | mg/dL | 3.9 ± 0.5 | 4.1 ± 0.7 | 3.8 ± 0.6 | 3.6 ± 0.4 | 4.8 ± 0.5 | 4.8 ± 0.3 | 4.3 ± 0.8 | 4.7 ± 0.6* |
| **METABOLIC** | | | | | | | | | |
| Glucose | mg/dL | 52.0 ± 15.1 | 56.0 ± 20.7 | 57.0 ± 20.4 | 46.3 ± 14.4 | 53.0 ± 10.7 | 57.2 ± 9.2 | 54.0 ± 11.6 | 51.5 ± 7.7 |
| Cholesterol | mg/dL | 333.0 ± 113.2 | 304.0 ± 91.1 | 328.7 ± 50.4 | 367.0 ± 69.9 | 251.3 ± 117.8 | 239.0 ± 117.1 | 229.2 ± 95.6 | 212.3 ± 114.1 |
| Triglycerides | mg/dL | 72.7 ± 28.3 | 66.3 ± 25.5 | 62.7 ± 1.5 | 66.0 ± 13.1 | 55.2 ± 23.3 | 54.5 ± 19.7 | 56.2 ± 25.4 | 52.2 ± 14.8 |
| Amylase | U/L | 146.0 ± 76.1 | 141.0 ± 60.8 | 140.3 ± 64.0 | 158.3 ± 67.0 | 147.2 ± 25.2 | 133.8 ± 21.9 | 133.2 ± 29.8 | 136.8 ± 22.8 |
| CPK | U/L | 259.3 ± 67.4 | 564.3 ± 255.5 | 339.3 ± 233.5 | 338.3 ± 300.8 | 850.2 ± 841.8[a] | 305.5 ± 214.8 | 1070.2 ± 1407.8[a] | 263.2 ± 128.6 |
| **ELECTROLYTES & MINERALS** | | | | | | | | | |
| Calcium | mg/dL | 9.1 ± 0.1 | 9.2 ± 0.1 | 9.4 ± 0.1 | 9.7 ± 0.1 | 9.2 ± 0.1 | 9.0 ± 0.1 | 9.3 ± 0.1 | 9.5 ± 0.4 |
| Magnesium | mEq/dL | 1.3 ± 0.1 | 1.2 ± 0.0 | 1.2 ± 0.1 | 1.3 ± 0.1 | 1.4 ± 0.2 | 1.3 ± 0.0* | 1.4 ± 0.0* | 1.4 ± 0.1 |
| Sodium | mEq/L | 148.7 ± 1.2 | 147.0 ± 1.7 | 147.0 ± 1.7 | 146.0 ± 2.0 | 147.2 ± 3.3 | 145.8 ± 2.2 | 147.5 ± 1.9 | 147.5 ± 2.1 |
| Potassium | mEq/L | 4.0 ± 0.5 | 3.9 ± 0.1 | 3.7 ± 0.1 | 4.0 ± 0.6 | 3.7 ± 0.4 | 3.6 ± 0.2 | 3.6 ± 0.2 | 3.7 ± 0.1 |
| Chloride | mEq/L | 109.3 ± 0.6 | 107.0 ± 1.0 | 107.7 ± 0.6 | 115.7 ± 15.3[b] | 106.0 ± 0.8** | 105.8 ± 1.0 | 108.0 ± 2.6 | 106.8 ± 3.3 |
| **HEMATOLOGY** | | | | | | | | | |
| RBC | $\times 10^6$/µL | 6.3 ± 0.3 | 6.1 ± 0.5 | 6.2 ± 0.3 | 6.3 ± 0.5 | 6.9 ± 0.2* | 6.4 ± 0.3 | 6.6 ± 0.2 | 6.8 ± 0.3 |
| HGB | g/dL | 13.0 ± 0.3 | 12.4 ± 0.7 | 12.9 ± 0.7 | 12.7 ± 0.4 | 13.8 ± 0.3* | 13.1 ± 0.5 | 13.6 ± 0.1 | 13.4 ± 0.1 |
| HCT | % | 44.3 ± 0.6 | 41.3 ± 1.2 | 42.0 ± 2.6 | 43.3 ± 2.1 | 48.0 ± 0.8*** | 43.5 ± 1.7 | 45.2 ± 1.3 | 45.8 ± 1.3 |
| MCV | fL | 71.0 ± 2.6 | 67.7 ± 4.0 | 68.3 ± 4.6 | 68.3 ± 3.8 | 69.8 ± 2.5 | 67.8 ± 4.2 | 69.0 ± 2.4 | 67.5 ± 1.3 |
| MCH | pg | 20.8 ± 0.5 | 20.4 ± 0.4 | 20.7 ± 0.7 | 20.1 ± 1.1 | 20.1 ± 0.4 | 20.4 ± 1.0 | 20.5 ± 0.6 | 19.7 ± 0.6 |

| | | | | | | | | | |
|---|---|---|---|---|---|---|---|---|---|
| MCHC | g/dL | 29.3 ± 0.6 | 30.3 ± 1.5 | 30.7 ± 1.2 | 29.3 ± 0.6 | 29.0 ± 0.8 | 30.2 ± 0.5 | 29.8 ± 0.5 | 29.2 ± 0.5 |
| WBC | ×10³/µL | 11.0 ± 1.7 | 11.1 ± 3.5 | 9.6 ± 2.4 | 10.6 ± 3.1 | 9.2 ± 0.6 | 8.5 ± 1.6 | 8.5 ± 2.1 | 9.5 ± 1.9 |
| Platelet count | ×10³/µL | 367.7 ± 38.0 | 398.3 ± 31.8 | 407.0 ± 58.6 | 433.0 ± 44.4 | 291.3 ± 40.1 | 355.0 ± 80.2 | 379.8 ± 83.5 | 396.0 ± 133.8 |
| Neutrophils | /µL | 7432 ± 2036 | 8108 ± 3165 | 6725 ± 2365 | 6912 ± 2016 | 5743 ± 445 | 6352 ± 1782 | 5648 ± 2393 | 5951 ± 2199 |
| Bands | /µL | 0 ± 0 | 0 ± 0 | 0 ± 0 | 0 ± 0 | 0 ± 0 | 0 ± 0 | 0 ± 0 | 0 ± 0 |
| Lymphocytes | /µL | 2930 ± 892 | 2281 ± 158 | 2349 ± 219 | 2623 ± 482 | 2713 ± 702 | 1609 ± 268** | 2260 ± 1152 | 2808 ± 1317 |
| Monocytes | /µL | 465 ± 85 | 538 ± 276 | 359 ± 65 | 933 ± 681 | 414 ± 60 | 272 ± 24 | 326 ± 18 | 443 ± 236 |
| Eosinophils | /µL | 172 ± 93 | 206 ± 78 | 167 ± 102 | 165 ± 224 | 330 ± 195 | 193 ± 150 | 176 ± 115 | 274 ± 212 |
| Basophils | /µL | 0 ± 0 | 0 ± 0 | 0 ± 0 | 0 ± 0 | 0 ± 0 | 0 ± 0 | 64 ± 129 | 0 ± 0 |

[a] Creatine kinase values in the anti-miR-128 arm at weeks 0 and 16 (850 ± 842 and 1,070 ± 1,408 U/L) are attributable to a single animal (UG228: 2,080 U/L pre-dose and 3,170 U/L at week 16), which returned to 402 U/L by week 24 on continued dosing. The remaining three treated animals ranged from 98 to 1,704 U/L across all timepoints. Transient creatine kinase elevation associated with restraint and ketamine sedation is well described in cynomolgus macaques.

[b] One saline animal (FR391) recorded a chloride of 133 mEq/L at week 24, against 105-109 mEq/L at its thirteen preceding draws and a concurrent sodium of 146 mEq/L. The value is reported as received from the analysing laboratory. The implied anion gap is not physiologically plausible and an analytical error is the most likely explanation; the value has been retained without correction and accounts for the elevated standard deviation shown.

### Supplementary Table 2. Serum Chemistry — DMD Pig Study

Data are presented as mean ± SD. N = 3 per group. ns, not significant; *p<0.05, **p<0.01, ***p<0.001 $DMD^{Y/-}$ Scramble vs. WT (the Anti-miR-128 group was not compared with WT); °p<0.05 Anti-miR-128 vs. Scramble (unpaired, two-tailed Welch t-test within timepoint / Δ W8−W0).

| Analyte | Units | Wild type (WT)[a] | | | $DMD^{Y/-}$ Scramble[a] | | | $DMD^{Y/-}$ Anti-miR-128[a] | | |
|---|---|---|---|---|---|---|---|---|---|---|
| | | Week 0 | Week 8 | Δ (W8 − W0) [%] | Week 0 | Week 8 | Δ (W8 − W0) [%] | Week 0 | Week 8 | Δ (W8 − W0) [%] |
| **MUSCLE DAMAGE** | | | | | | | | | | |
| CK | $10^3$ U/L | 1.18 ± 0.60 | 2.69 ± 1.68 | 267 ± 332 | 22.2 ± 11.7 | 103 ± 19.1* | 518 ± 343 | 37.3 ± 30.5 | 67.8 ± 26.2 | 160 ± 127 |
| ALAT | U/L | 23.3 ± 3.5 | 38.3 ± 5.5 | 65.5 ± 19.1 | 679 ± 206* | 511 ± 87.2* | -20.3 ± 21.0* | 627 ± 214 | 487 ± 83.9 | -19.0 ± 12.3 |
| ASAT | U/L | 41.7 ± 17.2 | 45.3 ± 9.7 | 19.2 ± 33.6 | 966 ± 717 | 840 ± 352 | 82.0 ± 184 | 1215 ± 742 | 1140 ± 420 | 16.6 ± 66.7 |
| LDH | U/L | 630 ± 87.4 | 603 ± 56.1 | -3.6 ± 7.9 | 3971 ± 667* | 8065 ± 2282* | 108 ± 64.5 | 4956 ± 1191 | 12434 ± 5152 | 164 ± 112 |
| **HEPATIC FUNCTION** | | | | | | | | | | |
| ALP | U/L | 481 ± 55.4 | 161 ± 19.1 | -66.4 ± 4.1 | 165 ± 11.6** | 114 ± 31.9 | -31.5 ± 13.3* | 154 ± 38.7 | 84.3 ± 9.7 | -43.2 ± 9.8 |
| Bilirubin, Total | µmol/L | 5.0 ± 1.8 | 0.7 ± 0.3 | -85.1 ± 6.5 | 2.0 ± 0.3 | 2.4 ± 1.0 | 14.4 ± 28.4* | 1.8 ± 0.9 | 2.4 ± 0.5 | 66.6 ± 69.8 |
| Total Protein | g/L | 27.1 ± 2.1 | 57.0 ± 3.8 | 112 ± 19.9 | 27.1 ± 2.9 | 45.5 ± 3.4* | 68.2 ± 7.0 | 23.1 ± 4.1 | 44.8 ± 2.3 | 98.3 ± 29.4 |
| Albumin | g/L | 23.6 ± 2.2 | 43.3 ± 1.2 | 84.4 ± 10.8 | 17.3 ± 2.8* | 32.4 ± 2.8* | 89.0 ± 13.1 | 14.0 ± 3.0 | 32.3 ± 1.8 | 137 ± 34.6 |
| GGT | U/L | 15.7 ± 4.0 | 24.7 ± 12.4 | 52.2 ± 31.5 | 27.3 ± 6.5 | 15.7 ± 2.9 | -38.8 ± 21.2* | 35 ± 16.8 | 22.3 ± 2.5° | -27.4 ± 22.6 |
| Amylase | U/L | 1553 ± 51.7 | 2218 ± 38.4 | 42.9 ± 5.6 | 1726 ± 178 | 1973 ± 403 | 13.6 ± 10.0* | 1452 ± 240 | 1917 ± 502 | 33.2 ± 25.5 |
| **RENAL FUNCTION** | | | | | | | | | | |
| BUN | mmol/L | 1.1 ± 0.0 | 3.5 ± 0.1 | 235 ± 13.3 | 0.6 ± 0.3 | 2.8 ± 1.7 | 492 ± 422 | 1.2 ± 0.4 | 2.6 ± 1.0 | 115 ± 20.9 |
| Creatinine | µmol/L | 48.7 ± 2.1 | 97.7 ± 1.5 | 101 ± 5.9 | 22 ± 3.5*** | 30.3 ± 12.7* | 34.4 ± 27.7* | 24 ± 7.0 | 35.3 ± 12.7 | 70.4 ± 97.4 |
| **METABOLIC** | | | | | | | | | | |
| Glucose | mmol/L | 5.5 ± 0.3 | 7.4 ± 0.4 | 35.6 ± 9.4 | 6.4 ± 1.1 | 6.0 ± 0.9 | -2.7 ± 28.3 | 5.2 ± 0.8 | 7.7 ± 3.2 | 54.4 ± 61.7 |
| Cholesterol | mmol/L | 2.9 ± 0.3 | 2.7 ± 0.2 | -6.4 ± 2.6 | 1.2 ± 0.3** | 1.7 ± 0.4* | 40.0 ± 9.8* | 1.2 ± 0.4 | 1.8 ± 0.2 | 67.7 ± 49.9 |
| Triglycerides | mmol/L | 0.8 ± 0.5 | 0.9 ± 0.2 | 49.3 ± 103 | 0.5 ± 0.1 | 0.8 ± 0.2 | 51.3 ± 46.1 | 0.6 ± 0.2 | 0.9 ± 0.3 | 68.8 ± 48.5 |
| Fructosamine | µmol/L | 131 ± 8.7 | 209 ± 5.7 | 59.2 ± 5.0 | 134 ± 3.5 | 180 ± 11.1* | 34.9 ± 5.8* | 124 ± 22.2 | 181 ± 11.9 | 50.5 ± 30.6 |
| Lactate | mmol/L | 2.3 ± 0.4 | 2.1 ± 0.1 | -9.2 ± 10.0 | 2.7 ± 0.9 | 3.9 ± 0.6* | 57.0 ± 53.0 | 1.5 ± 0.1 | 3.2 ± 0.8 | 112 ± 44.5 |
| Lipase | U/L | 6.5 ± 0.8 | 8.5 ± 0.8 | 31.6 ± 10.2 | 4.7 ± 0.7* | 5.3 ± 0.4* | 14.2 ± 6.2 | 4.3 ± 0.7 | 5.8 ± 0.3 | 38.5 ± 21.9 |
| **ELECTROLYTES & MINERALS[b]** | | | | | | | | | | |
| Phosphate | mmol/L | 2.2 ± 0.3 | 3.2 ± 0.4 | 44.4 ± 9.1 | 2.2 ± 0.2 | 3.4 ± 0.3 | 53.8 ± 12.8 | 2.0 ± 0.4 | 3.5 ± 0.3 | 72.7 ± 15.5 |
| Calcium | mmol/L | 2.1 ± 0.0 | 2.8 ± 0.2 | 31.9 ± 6.8 | 2.3 ± 0.1 | 2.3 ± 0.1* | 1.4 ± 3.1* | 2.1 ± 0.3 | 2.4 ± 0.2 | 19.0 ± 8.8 |
| Magnesium | mmol/L | 0.8 ± 0.1 | 1.0 ± 0.1 | 37.1 ± 15.7 | 0.9 ± 0.1 | 0.9 ± 0.1 | 0.1 ± 13.0* | 0.8 ± 0.1 | 0.9 ± 0.0 | 14.1 ± 13.4 |

[a] WT and $DMD^{Y/-}$ pigs in the different treatment groups were littermates. Reference ranges for pigs are not given as published porcine intervals vary by breed and age.

[b] Sodium, chloride, and potassium concentrations were consistently below the porcine reference ranges, indicating a possible analytical or preanalytical issue. These values were therefore excluded from further consideration.

### Supplementary Table 3. RT-qPCR Primers.

Primer sequences used for RT-qPCR gene expression analysis.

| Gene Symbol | Species | Forward Oligo (5'→3') | Reverse Oligo (5'→3') |
|---|---|---|---|
| *Ppargc1a* | Mouse | TATGGAGTGACATAGAGTGTGCT | CCACTTCAATCCACCCAGAAAG |
| *Tfam* | Mouse | TGGCAGTCCATAGGCACCGTATT | ACAGACAAGACTGATAGACGAGGG |
| *Atp5g* | Mouse | CCTGTGCCTGTCTTTCTACC | CCTTCCACACTCTGCCTTATC |
| *Sod2* | Mouse | CAGGATGCCGCTCCGTTAT | TGAGGTTTACACGACCGCTG |
| *Ppara* | Mouse | GTGCAGCCTCAGCCAAGTT | TGGGGAGAGAGGACAGATGG |
| *Drp1* | Mouse | GTTCCACGCCAACAGAATAC | CCTAACCCCCTGAATGAAGT |
| *Fis1* | Mouse | AAGTATGTGCGAGGGCTGT | TGCCTACCAGTCCATCTTTC |
| *Mfn1* | Mouse | GACCCGTGCGAAAGAGAGAG | TCGAGCAAAAGTAGTGGCCA |
| *Mfn2* | Mouse | ATGTTACCACGGAGCTGGAC | AACTGCTTCTCCGTCTGCAT |
| *Opa1* | Mouse | ATACTGGGATCTGCTGTTGG | AAGTCAGGCACAATCCACTT |
| *R18s* | Mouse | GCAATTATTCCCCATGAACG | GGCCTCACTAAACCATCCAA |
| *Tnfa* | Mouse | CCCTCACACTCAGATCATCTTCT | GCTACGACGTGGGCTACAG |
| *Prkaa2* | Mouse | GGAGAACACCAATTGACAGGC | GTGCTGATCACCTGGTAGAGT |
| *Sirt1* | Mouse | CGGCTACCGAGGTCCATATAC | CGCTTTGGTGGTTCTGAAAGG |
| *Mt-co3* | Mouse | AAATTCCTGTTGGAGGTCAGCAGC | TCGTACCACAGGCATTGTGATGGA |
| *Actb* | Mouse | TGTGACTTTGTGGTGTGGCTG | CACCTGCACTCTGGGTAAGGA |

### Supplementary Table 4. Trait Label Mapping for GWAS Credible Set Analysis.

Mapping of GWAS Catalog reported trait strings to simplified labels and broad trait categories used throughout the text and figures. Traits are grouped by category.

| Reported Trait (GWAS Catalog) | Label (this study) |
|---|---|
| **Anthropometric** | |
| Body mass index | BMI |
| Obesity | Obesity |
| Obesity due to excess calories | Obesity |
| Morbid obesity (PheCode 278.11) | Morbid obesity |
| Body fat percentage (UKB data field 23099) | Body fat % |
| Waist-hip ratio | Waist-hip ratio |
| Body mass index (BMI) (UKB data field 21001) | BMI |
| Body mass index (BMI) (UKB data field 23104) | BMI |
| Body mass index (UKB data field 21001) | BMI |
| Arm fat mass left (UKB data field 23124) | Arm fat mass |
| Arm fat mass right (UKB data field 23120) | Arm fat mass |
| Arm fat percentage left (UKB data field 23123) | Arm fat % |
| Arm fat percentage right (UKB data field 23119) | Arm fat % |
| Waist circumference (UKB data field 48) | Waist circumference |
| Whole body fat mass (UKB data field 23100) | Whole body fat mass |
| Trunk fat percentage (UKB data field 23127) | Trunk fat % |
| Hip circumference (UKB data field 49) | Hip circumference |
| **Lipids** | |
| HDL cholesterol | HDL |
| HDL cholesterol levels | HDL |
| HDL cholesterol levels (UKB data field 30760) | HDL |
| High density lipoprotein cholesterol levels | HDL |
| High density lipoprotein cholesterol levels (UKB data field 30760) | HDL |
| High-density lipoprotein levels (MTAG) | HDL |
| Low density lipoprotein cholesterol levels | LDL |
| LDL direct levels (UKB data field 30780) | LDL |
| Cholesterol levels | Total cholesterol |
| Cholesterol levels (UKB data field 30690) | Total cholesterol |
| Total cholesterol levels | Total cholesterol |
| Total esterified cholesterol levels | Total cholesterol |
| Non-HDL cholesterol levels | Non-HDL cholesterol |
| Apolipoprotein B levels | ApoB |
| Apolipoprotein B levels (UKB data field 30640) | ApoB |
| Apolipoprotein A1 levels | ApoA1 |
| Phospholipids in large HDL | Phospholipids (HDL) |
| Phospholipids in very large HDL | Phospholipids (HDL) |
| Phospholipids to Total Lipids in Large HDL percentage | Phospholipids/total lipids ratio (HDL) |
| Phospholipids to Total Lipids in Medium HDL percentage | Phospholipids/total lipids ratio (HDL) |
| Phospholipids to Total Lipids in Medium HDL percentage (PGS-adjusted) | Phospholipids/total lipids ratio (HDL) |
| Phospholipids to total lipids ratio in medium HDL | Phospholipids/total lipids ratio (HDL) |

| | |
|---|---|
| Phospholipids to Total Lipids in Large LDL percentage | Phospholipids/total lipids ratio (LDL) |
| Phospholipids to Total Lipids in Large LDL percentage (PGS-adjusted) | Phospholipids/total lipids ratio (LDL) |
| Omega-3 fatty acids | Omega-3 FA |
| Omega-3 fatty acids levels | Omega-3 FA |
| Omega-3 fatty acid levels | Omega-3 FA |
| Docosahexaenoic acid levels | DHA |
| Circulating docosahexaenoic acid levels | DHA |
| Sphingomyelins levels | Sphingomyelins |
| **Metabolic** | |
| Urate levels | Urate |
| Urate levels (UKB data field 30880) | Urate |
| Frailty (Factor 4 - Metabolic Problems) | Frailty (metabolic) |
| Polyneuropathy in diabetes (PheCode 250.6) | Diabetic polyneuropathy |
| **Cardiovascular event** | |
| Sudden cardiac arrest in coronary artery disease | Cardiac arrest (in CAD) |
